# All-atom Simulations Uncover the Molecular Terms of NKCC1 Transport Mechanism

**DOI:** 10.1101/2021.05.12.443869

**Authors:** Pavel Janoš, Alessandra Magistrato

## Abstract

The secondary-active Na-K-Cl Cotransporter 1 (NKCC1), member of the Cation Chloride Cotransporters (CCC) family, ensures the electroneutral movement of Cl^-^, Na^+^, K^+^ ions across cellular membranes. NKCC1 regulates Cl^-^ homeostasis and cell volume, handling a pivotal role in transepithelial water transport and neuronal excitability. Aberrant NKCC1 transport is hence implicated in a variety of human diseases (hypertension, renal disorders, neuropathies, cancer). Building on the newly-resolved NKCC1 cryo-EM structure, all-atom enhanced sampling simulations unprecedentedly unlock the mechanism of NKCC1-mediated ions transport, assessing the order and the molecular basis of its interdependent ions translocation. Our outcomes strikingly advance the understanding of the physiological mechanism of CCCs transporters and disclose a key role of CCC-conserved asparagine residues, whose side-chain promiscuity ensures the transport of both negatively and positively charged ions along the same translocation route. This study sets a conceptual basis to devise NKCC-selective inhibitors to treat diseases linked to Cl^-^ dishomeostasis.

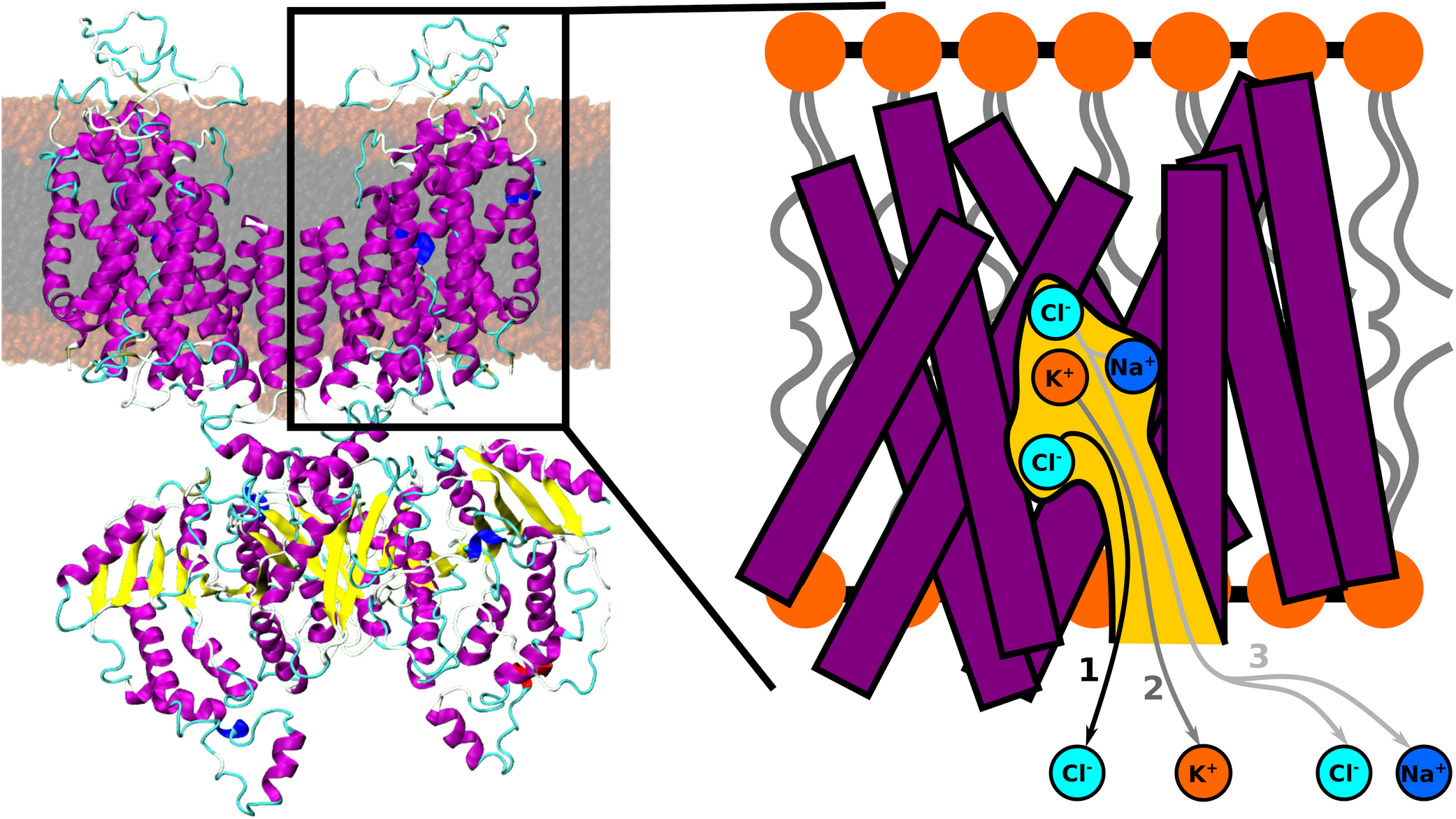

## Introduction

The sodium potassium chloride cotransporters (NKCC), part of the cation-chloride cotransporter (CCC) family, harness the K^+^ and/or Na^+^ concentration gradient, generated by the Na^+^-K^+^-ATPase,^1–3^ to uptake/extrude Cl^-^ ions in/out the plasma membrane of distinct cell types.^3,4^ Among CCCs, two are Na^+^ dependent (NKCC1 and NKCC2), and four are Na^+^ independent (KCC1-4).

NKCCs exploit the energy of the Na^+^ gradient to accumulate cellular Cl^-^ beyond their electro-chemical equilibrium, while KCCs take advantage of the outward K^+^ gradient to extrude Cl^-^ ions from the cell. In many cell types, NKCC(s) and KCC(s) are reciprocally regulated by kinases and phosphatases, and this tight control is critical to maintain intracellular Cl^-^ homeostasis and to regulate cell volume. Among CCCs, NKCC1 is the most extensively characterized. By mediating Cl^-^ uptake, NKCC1 triggers a concomitant obligatory water influx as major defense mechanism against cell shrinkage.^5^

NKCC1 is widely distributed throughout the human body, being present (i) in the basolateral membrane of exocrine glands, where it uptakes ions from the blood, thus being critical for salt re-absorption; (ii) in the endolymph fluid present in the inner part of ear, playing a key role for hearing, (iii) in neurons where, coadjuvated by the opposing action of KCC2, it regulates neuronal excitability and γ-aminobutyric acid (GABA)-mediated signaling.

The manifold centrality of NKCC1 transporter is overwhelmingly evidenced by its implications in neonatal seizures, neuropathic pain, and neurological disorders (including Down syndrome and autism spectrum disorders),^6–8^ also in hypertension^9^ and renal disturbances (Gitelman’s, Bartter’s and Andermann’s syndromes).^10^ Withal, a growing body of evidences links NKCC1 expression to tumor infiltration and progression in multiple cancer types (lung adenocarcinoma, prostate cancer, esophageal squamous carcinoma, brain tumors).^11^ In this scenario, NKCC1 represents an appealing pharmacological target to treat a variety of diseases associated with Cl^-^ dis-homeostasis. Indeed, marketed drugs targeting NKCC1, such as the loop diuretics furosemide and bumetanide, which are commonly prescribed for pulmonary edema,^12^ also exhibit anti-seizure,^13^ anti-epileptic,^14^ antiparkinsonian^15^ and anti-tumor activity.^11^ Nevertheless, a poor selectivity jeopardizes their prolonged use for chronic patients.^16^

The NKCC1 structure exhibits a pseudo-symmetric topology composed by two inverted repeats of five transmembrane helices forming a central binding cavity. NKCC1 transports the Na^+^, K^+^ and Cl^-^ ions in a 1:1:2 stoichiometry^17^ across cellular membranes via a so-called alternating-access mechanism whereby the transporter visits outward-facing, occluded, and inward-facing conformation.^16,18,19^ Specifically, NKCC1 presents a homodimeric transmembrane domain, and N-terminal and C-terminal domains that contribute to dimer assembly and possibly also modulate ions trafficking. ^20,21^

A recent cryo-EM structure of dimeric zebrafish NKCC1^20^ grasped the transporter in its inward-facing state (Figure 1), also tracing two Cl^-^ and one K^+^ ions in their binding sites. The position of the Na^+^ binding site was, instead, inferred based on structural homology with other transporters. Although this structure provided ground-breaking information to foresee the NKCC1 function, many facets of its transport mechanism remain yet to be resolved.

**Figure 1.**
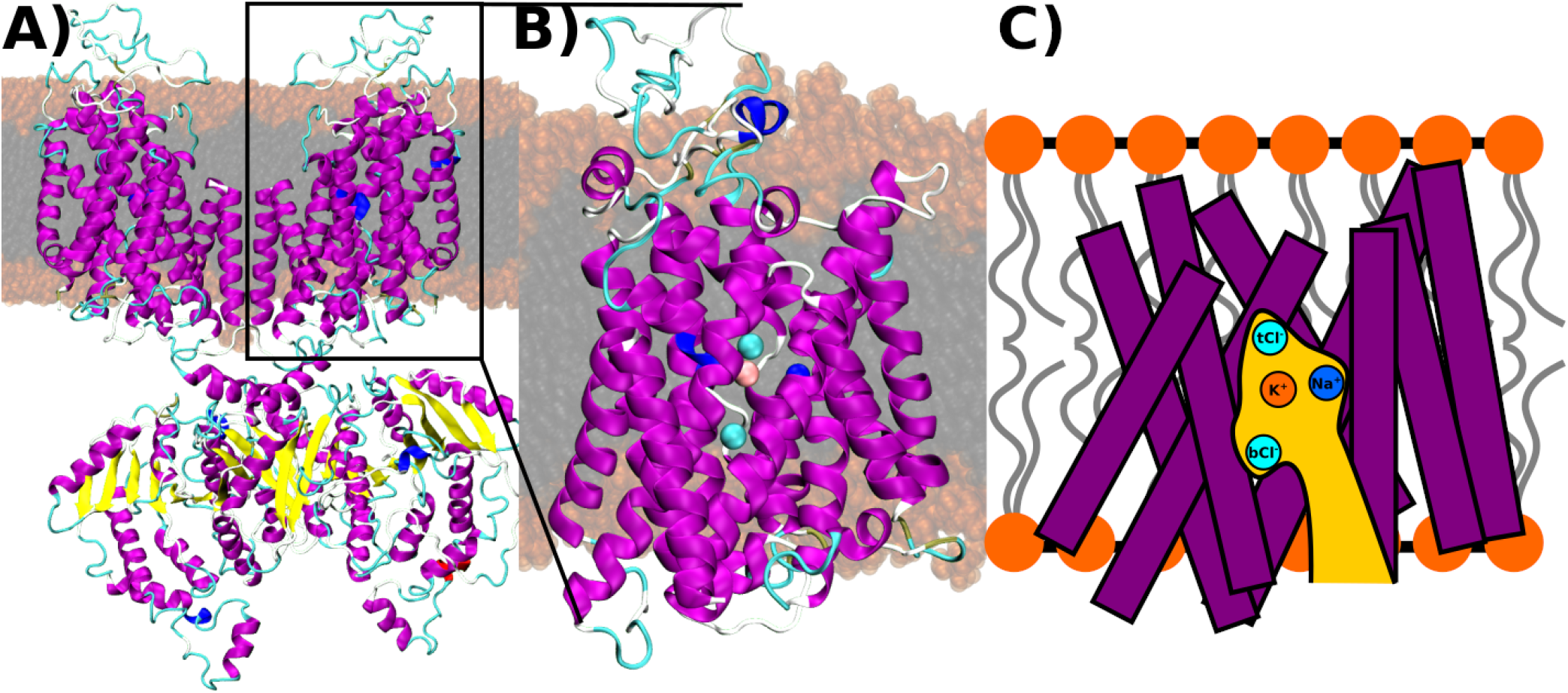
The NKCC1 transporter (A). The model used to assess the in-cell transport mechanism of all ions (B). NKCC1 is depicted as magenta and yellow new-cartoons for helix and β-sheet structures, respectively. The Cl^-^, Na^+^ and K^+^ ions are shown as cyan, blue, and orange van deer Waals spheres, respectively. The hydrophilic head groups and hydrophobic tails of the 1-palmitoil-2-oleoyl-sn-glycero-3-phosphocholine (POPC) lipids are depicted in orange and gray surfaces, respectively. (C) Sketch of the reduced NKCC1 model with the helices depicted as magenta rectangles, the binding cavity highlighted in yellow and the Cl^-^, Na^+^ and K^+^ ions sketched in cyan, blue and orange circles, respectively.

In this study, we employed all-atom enhanced sampling simulations to unprecedentedly and exhaustively assess the NKCC1-aided ions transport mechanism, picturing the free energy surface of whole process, tracing the most energetically-viable translocation routes, outlining the relative order of in-cell ions release and the molecular basis of their cooperative transport. A thorough understanding of NKCC1 function advances our knowledge on the whole CCC family, setting a conceptual basis to design selective inhibitors of this pivotal, yet emerging, drug-target.

## Materials and Methods

### Model Building

Two NKCC1 models were built based on the cryo-EM structure (PDB id: 6NPL) from zebrafish^20^: In the first (large) NKCC1 model we omitted the first 205 N-term residues missing in the cryo-EM structure, while other missing loops were modeled with the MODELLER program.^22^ The second (small) model NKCC1_cut_ encompassed only a monomer of transmembrane domain (residues 206-677). The first model was used as a reference to assess the structural stability of the NKCC1_cut_, which was exploited to investigate the ions transport mechanism. The Protein Preparation Wizard of Schrodinger ^23^ was used to assign the protonation states of ionizable residues. The positions of bound K^+^ and Cl^-^ were taken from the cryo-EM structure, while the Na^+^ ion was manually placed into the binding site proposed in ref.^20^ on the basis of structural homology with other Na^+^ transporters. Both models were inserted into a pre-equilibrated 1-palmitoil-2-oleoyl-sn-glycero-3-phosphocholine (POPC) membrane bilayer using CHARMM-GUI.^24,25^ For the large NKCC1 model the size of the membrane was 160 Å x 160 Å and contained 335 and 325 POPC molecules in upper and lower leaflet, respectively. Three POPC lipids, present in the dimerization interface in the cryoEM structure, were retained in the model. The system was hydrated in the z direction by adding a shell of 22.5 Å of water in the z direction leading to 469231 atoms. The NKCC1cut model, instead, was inserted into a membrane of 110 Å x 110 Å size, which contained 158 and 152 lipids in upper and lower leaflet, respectively. After the addition of a 22.5 Å layer of water in the z direction, the total number of atoms was 139140. The system was described using Amber FF14SB^26^ and Lipid17^27^ forcefields for the protein and POPC lipids, respectively; and was solvated using the TIP3P water model^28^ with 150 nM concentration of NaCl. Joung and Chetham parameters were used for all the Na^+^, Cl^-^ and K^+^ ions.^29^

### Molecular dynamics simulations details

Molecular dynamics (MD) simulations were performed using GROMACS 2020 simulations package^30,31^ coupled with PLUMED 2.7.0 for metadynamics (MTD) simulations.^32,33^ The systems were equilibrated in 8 steps with an increasing MD simulations length (0.5 ns, 0.25 ns, 0.5 ns, 0.5 ns, 0.5 ns, 0.5 ns, 0.5 ns, 2.0 ns) and decreasing restraints on the protein (5000 kJ/mol·nm, 4000 kJ/ mol·nm, 3000 kJ/mol·nm, 2000 kJ/mol·nm, 1000 kJ/mol·nm, 500 kJ/mol·nm, 250 kJ/mol·nm, 50 kJ/ mol·nm, 0 kJ/mol·nm) and the membrane (2000 kJ/mol·nm, 2000 kJ/mol·nm, 1000 kJ/mol·nm, 500 kJ/mol·nm, 250 kJ/mol·nm, 100 kJ/mol·nm, 50 kJ/mol·nm, 0 kJ/mol·nm, 0 kJ/mol·nm). Restraints on Cα atoms with 50 kJ/mol·nm were applied in the last step. The Cα restraints were kept for the first 100 ns of MD simulations to allow for membrane equilibration. The simulations were then extended up to 500 ns without restraints. Hydrogen mass repartitioning was used for the large NKCC1 model system to allow a 4 fs time-step.^34^ For all other simulations a time-step of 2 fs time was used.

### Metadynamics simulations details

The order of ions dissociation was determined using MTD simulations biasing all four ions using, as collective variables (CVs), the distance of each ion to its binding sites. The parameters employed in these initial MTD simulations were: deposition time = 10 ns, hill height = 0.42 kJ/mol, σ = 0.3 Å. Once an ion dissociated, the simulation was restarted biasing the dissociation of the remaining ions until the dissociation of all ions was achieved. The simulations leading to the dissociation of each ion were repeated three times to confirm the non-stochastic nature of the observed series of dissociation events. Detailed transport mechanisms of each ion were obtained from well-tempered (WT) volumetric-based MTD simulations.^35^ This approach employs spherical coordinates of the ligand (ρ, θ, φ) as CVs for the WT-MTD simulations. The ρ CV represents the distance of the ligand from the center of the coordinate system, which here corresponds to the center of mass of the core helices (H1, H2, H3, H6, H7, H8, H10). The θ and φ CVs define angles accounting for the orientation of the ligand within a sphere of radius ρ. The sampling space is limited by restraining the ρ coordinate at 18 Å, which corresponds to the distance, where the ion egresses the channel surface. Nevertheless, a restraint at this ρ value does not allow the full ion dissociation into the bulk water, thus preventing the exchange between the bound ion with those present in the bulk water, which would hinder the sampling of multiple binding/unbinding events and compromise the convergence of the MTD simulation. When assessing the Na^+^ translocation mechanism, an additional restraint was needed to prevent the entrance into the channel of the bulk Cl^-^ ions, which could interfere with the Na^+^ dissociation process. To this end, we restrained to 0.1 the value of the coordination number of the solvated Cl^-^ ions with respect to the Cl^-^ binding sites. In these WT-MTD simulations we used a deposition time = 5 ns, hill height = 1.0 kJ/mol, σ_rho_ = 0.5 Å, σ_theta_ = π/16, σ_phi_ = π/8, and bias factor = 10, following a previously applied protocol ^35^. The simulation for the dissociation of each ion was run for 200 ns.

For analyses purposes the free energy profiles were reweighted onto the distance from the center, ρ, as CV1, and the coordination number of the ion with respect to protein heavy atoms, as CV2. The coordination number uses formula: 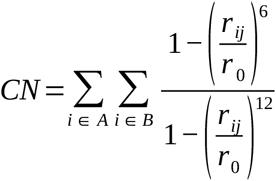 where A is the set of non-hydrogen atoms of the ligand, B the set of non-hydrogen atoms of the protein and the r0 is a threshold to define a formed contact. We use different r0 for different ions to account for their different van der Waals diameters: r_0Cl_ = 4 Å, r_0K_ = 3.5 Å, r_0Na_ = 3.5 Å.

Due to the observed cooperativity during dissociation of the Na^+^ and top (inner-most protein bound) tCl^-^ ions, these were biased simultaneously using two CVs accounting for the distance of each ion from its respective binding pocket. To this end we used two CVs, accounting for the distance of each ion from its respective binding pocket (Table S1). In this 400 ns-long WT-MTD we used hill deposition time = 5 ns, hill height = 2.0 kJ/mol, σ = 0.25 Å, bias factor = 5. Restraining wall was placed at 20 Å for both CVs to prevent the full dissociation of the ions into the bulk solvent.

In all MTD simulations the error of the free energy profiles was calculated as the standard deviation of different time averages from different simulation blocks.

## Results

### Relative Order of Ions Translocation

We initially validated the structural stability of a reduced transmembrane NKCC1 model, as compared to a complete one (Supplementary Information Figures S1-S3), by performing extensive Molecular Dynamics (MD) simulations of NKCC1 embedded in a 1-palmitoil-2-oleoyl-sn-glycero-3-phosphocholine (POPC) membrane bilayer. Our model, reliably reproduces the experimentally proposed ion binding sites (Figures S4-S6).

Aiming to unravel the order of ions translocations from the fully ions-loaded inward-facing NKCC1, we performed preliminary metadynamics (MTD) simulations in which we induced (biased) simultaneously the departure of all ions (Figures S7-S9). As a result, we first observed the most cytosol-exposed (bottom, b) Cl^-^ dissociation. Its departure affected the integrity of the K^+^ binding site, triggering its consequent release. Conversely, it was not possible to unequivocally discriminate relative dissociation order of the Na^+^ and the most deeply buried (top, t) Cl^-^ ions since distinct MTD simulations prompted either their concomitant translocation or the dissociation of tCl^-^ closely followed by that of Na^+^ (Figure S9).

After assessing the sequence of in-cell ions transport events, we performed ion-focused-MTD simulations exploring in more detail the dissociation mechanism of each ion, while also appraising their underlying kinetic and thermodynamic profile.

### Transport Mechanism of the Most Cytosol-Oriented Cl^-^ Ion

We initially characterized the free energy surface (FES) of the bCl^-^ ion translocation (Figure 2) since it was the first ion to exit the channel in preliminary MTD simulations. Well-tempered-MTD simulations disclosed that bCl^-^ moves from its initial binding site (**0**_**bCl**_) to the first metastable minimum, **1**_**bCl**_, overcoming a free energy barrier ΔG^‡^_0→ 1_ of 1.6 ± 0.6 kcal/mol, where it loses the coordination with the backbone amide hydrogens of residues Gly421 and Asn220, and with the hydroxyl hydrogen of Tyr611.

**Figure 2.**
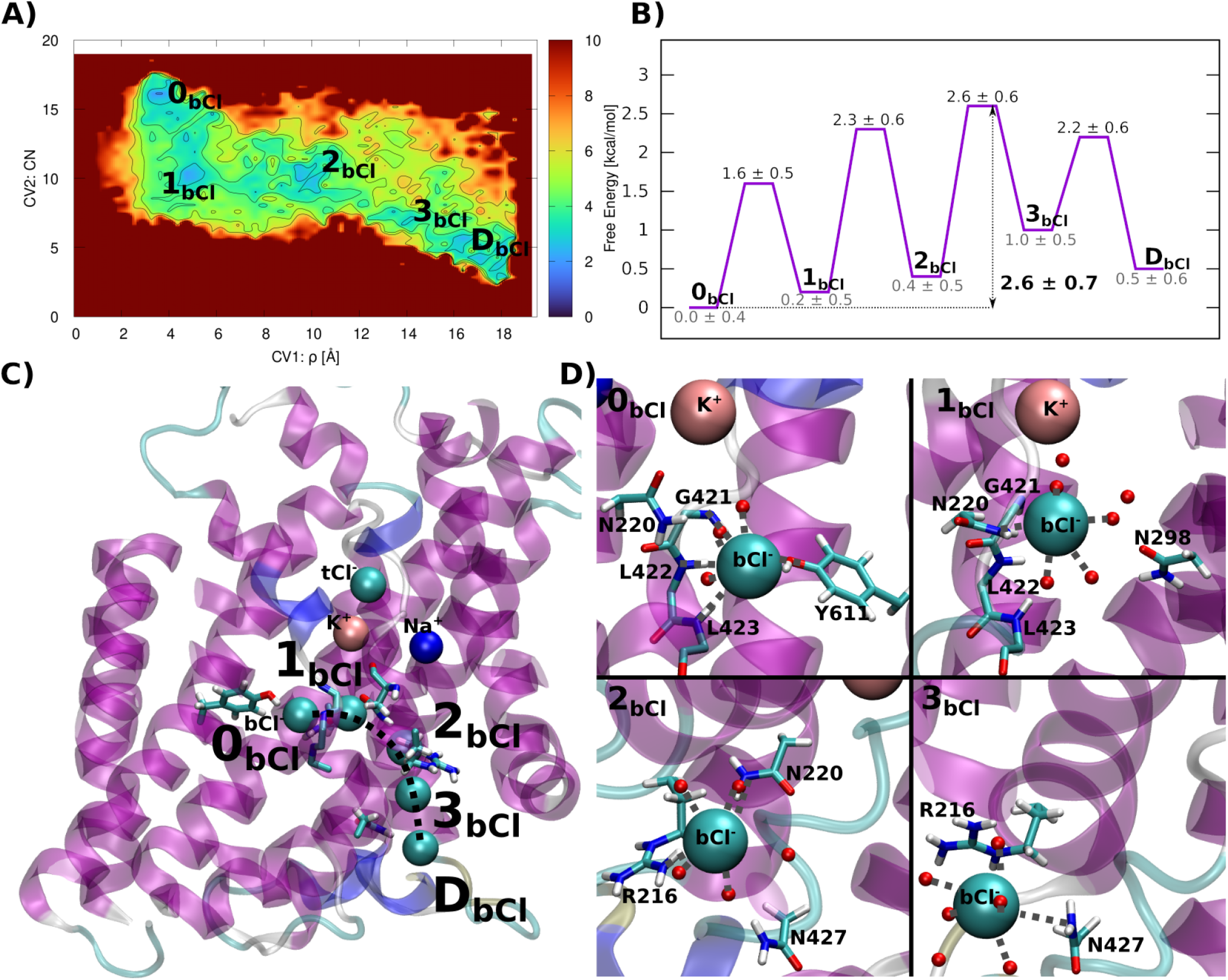
Transport of the most cytosol-exposed (bottom, b) Cl^-^ ion. (A) Free energy surface (kcal/mol) as a function of Collective Variable (CV) 1 and CV2 defined as the distance (Å) of bCl^-^ from the protein center of mass and of its coordination number to the protein residues, respectively. The free energy isosurface (kcal/mol) is shown from blue to red color; (B) Simplified graph showing the free energy of minima and transition states visited by bCl^-^ along its translocation route (relative free energies are reported with respect to the most stable minimum); (C) View of the bCl^-^ ion transport pathway with the consecutive positions of the ion highlighted with a dashed line. The positions of the top (t) Cl^-^, Na^+^ and K^+^ ions are also depicted. (D) Close-up views of the initial binding site and the most relevant states visited by bCl^-^ along its dissociation route. The coordination sphere of the ion is highlighted with gray dashed lines. The NKCC1 protein is depicted as magenta new cartoons, the Cl^-^, Na^+^ and K^+^ ions as cyan, blue and pink van der Waals spheres, respectively. The residues and water molecules are shown in licorice and balls (without hydrogens), respectively.

We remark that all ions are hexacoordinated through-out our simulations, with water molecules completing the coordinate sphere in the absence of suitable protein residues. For clarity, water molecules are omitted from the description of the ion coordination spheres here and in the following sections. In **1**_**bCl**_ the bCl^-^ is closer to K^+^, being stabilized by direct and water-mediated interactions with residues Asn220 and Asn298, respectively. Next, surmounting a ΔG^‡^_1→ 2_ = 2.1 ± 0.8 kcal/mol, bCl^-^ heads to a second minimum **2**_**bCl**_, where it is caught by Arg216 and Asn220 sidechains, and by Asn427 side-chain through a water-mediated interaction. Remarkably, **2**_**bCl**_ corresponds to a low-affinity Cl^-^ binding site discussed in ref.^20^, which was proposed to be a transient minimum of CLC transporters.^36^ Hereafter, bCl^-^ moves into the exit part of the channel, and, after dissociating from Asn220 at the cost of ΔG^‡^_2→ 3 **bCl**_ = 2.2 ± 0.8 kcal/mol, it reaches a third minimum **3** where it is bound by Asn427 side-chain amide hydrogen. Ultimately, disengaging from Asn427 (ΔG^‡^_3→ D dbCl_ = 1.2 ± 0.8 kcal/mol), bCl^-^ becomes fully solvated. The overall ΔG^‡^ of the bCl^-^ ion is of 2.6 ± 0.7 kcal/mol, and the multiple metastable minima of the FES lie 0.5-1 k/mol above the main (**0**_**bCl**_) minimum.

### Transport Mechanism of the K^+^ Ion

The dissociation of K^+^ encompasses four intermediate-states **1**_**K**_**-4**_**K**_ (Figure 3, S11). The Thr420 residue, coordinating K^+^ via its backbone carbonyl oxygen in the initial binding site (**0**_**K**_), is linked to Gly421, whose amide group forms, in turn, the bCl^-^ ion coordination sphere. Therefore, after bCl^-^ dissociation has occurred, Thr420 undergoes a conformational remodeling, destabilizing the K^+^ binding site. Next, K^+^ progresses along its in-cell delivery route by dissociating from the backbone carbonyl oxygen of Ala418 (ΔG^‡^_0→ 1_ = 0.7 ± 0.5 kcal/mol), thus heading to minimum **1**_**K**_. Here, the K^+^ is bound by the Asn220 sidechain and the carbonyl oxygens of Asn220 and Thr420. By surmounting a ΔG^‡^_1→ 2_ = 1.9 ± 0.5 kcal/mol, K^+^ moves to **2**_**K**_, where it is engaged by the carbonyl oxygen of Asn298 and by water-mediated interaction with Asn220 side-chain (Figure 3). From **2**_**K**_, K^+^ reaches **3**_**K**_ (ΔG^‡^_2→ 3_ = 4.1 ± 0.7 kcal/mol), where it is coordinated by Glu353 carboxyl. Finally, the K^+^ encounters the slowest step of its entire dissociation route (ΔG^‡^_3→ 4_ = 4.8 ± 0.7 kcal/mol), where, after losing its coordination to Glu353 side-chain, it reaches the final metastable state **4**_**K**_. Here, K^+^ is coordinated by sidechain oxygen of Asn427. The last dissociation step occurs at a modest free energy cost (ΔG^‡^_4→ D_ = 1.0 ± 0.6 kcal/mol). As a result, the overall ΔG^‡^_dK_ for K^+^ egress is 5.8 ± 0.5 kcal/mol and the metastable states visited along the translocation paths lie about 1-1.5 kcal/mol higher free energy as compared to **0**_**K**_.

**Figure 3.**
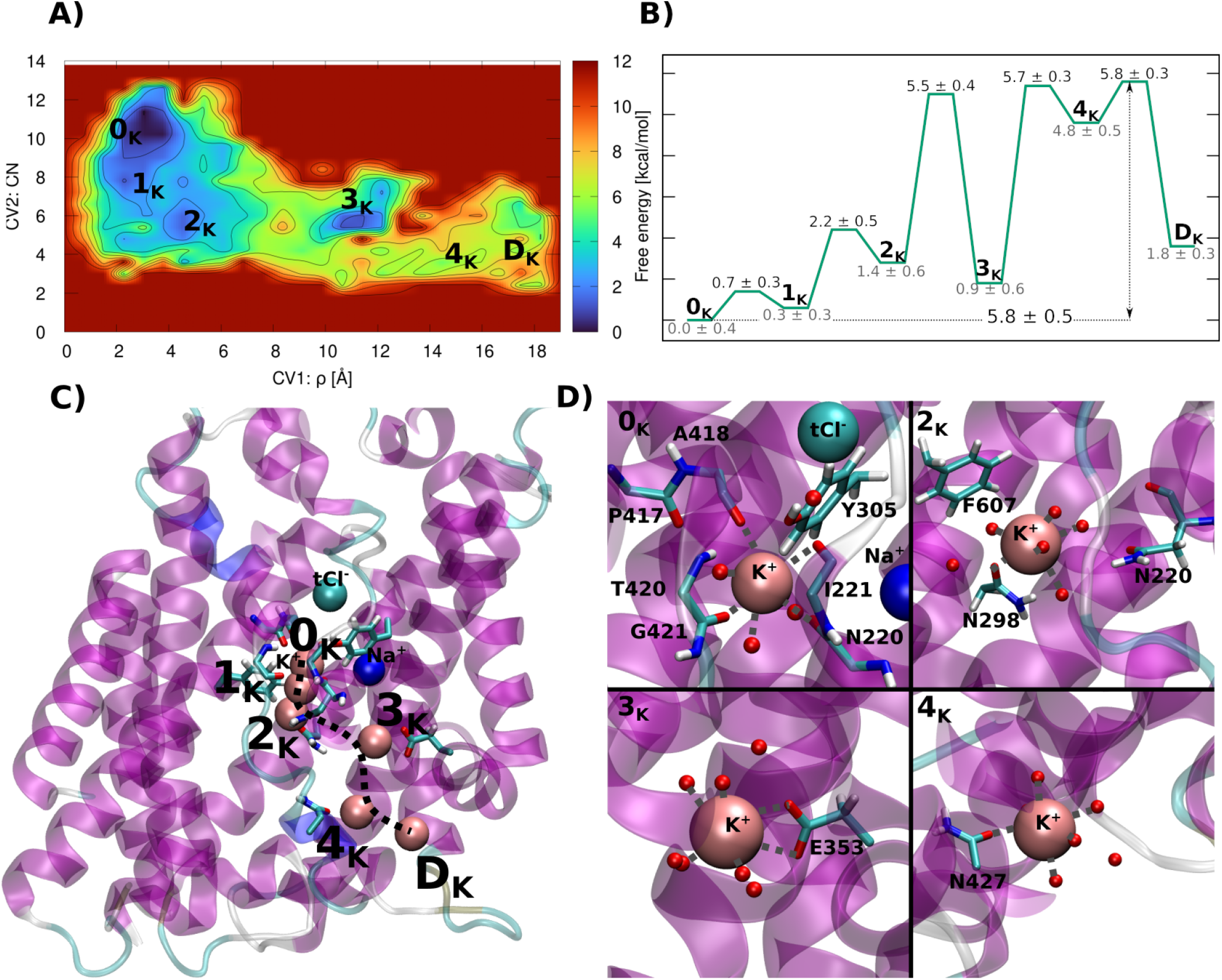
(A) Free energy surface (kcal/mol) reported as a function of Collective Variable (CV) 1 and CV2 defined as the distance (Å) of K^+^ from the protein center of mass and its coordination number to the protein residues, respectively. The free energy isosurfaces are shown from blue to red color; (B) Simplified graph showing the free energy of minima and transition states met along the translocation path (free energies are reported relative to the main minimum) (C) Image of the K^+^ ion dissociation steps sketched on the NKCC1 channel with the subsequent positions of the ion highlighted with a black dashed line. The positions of the top (t)Cl^-^ and Na^+^ ions are also depicted. (D) Close-up view of the initial binding site and selected relevant minima of the K^+^ ion dissociation process. The K^+^ coordination sphere is highlighted with gray dashed lines. The NKCC1 protein is depicted as magenta new cartoons, the Na^+^ and K^+^ ions as blue and pink van der Waals spheres, respectively. Protein residues and water molecules lining the minima along the translocation pathway are shown in licorice and balls (without hydrogens), respectively.

### Transport Mechanism of the Most NKCC1-Buried Cl^-^ Ion and/or the Na^+^ Ion

As detailed above, tCl^-^ and Na^+^ ions cooperatively dissociate from NKCC1. Thus, we unlocked their egress mechanism by concomitantly biasing the distance of each ion from its initial binding site. This led to an intricate FES, where the dissociation of either the tCl^-^ ion or both tCl^-^ and Na^+^ ions proceeded via three entangled paths (Paths 1-3, Figure 4).

**Figure 4.**
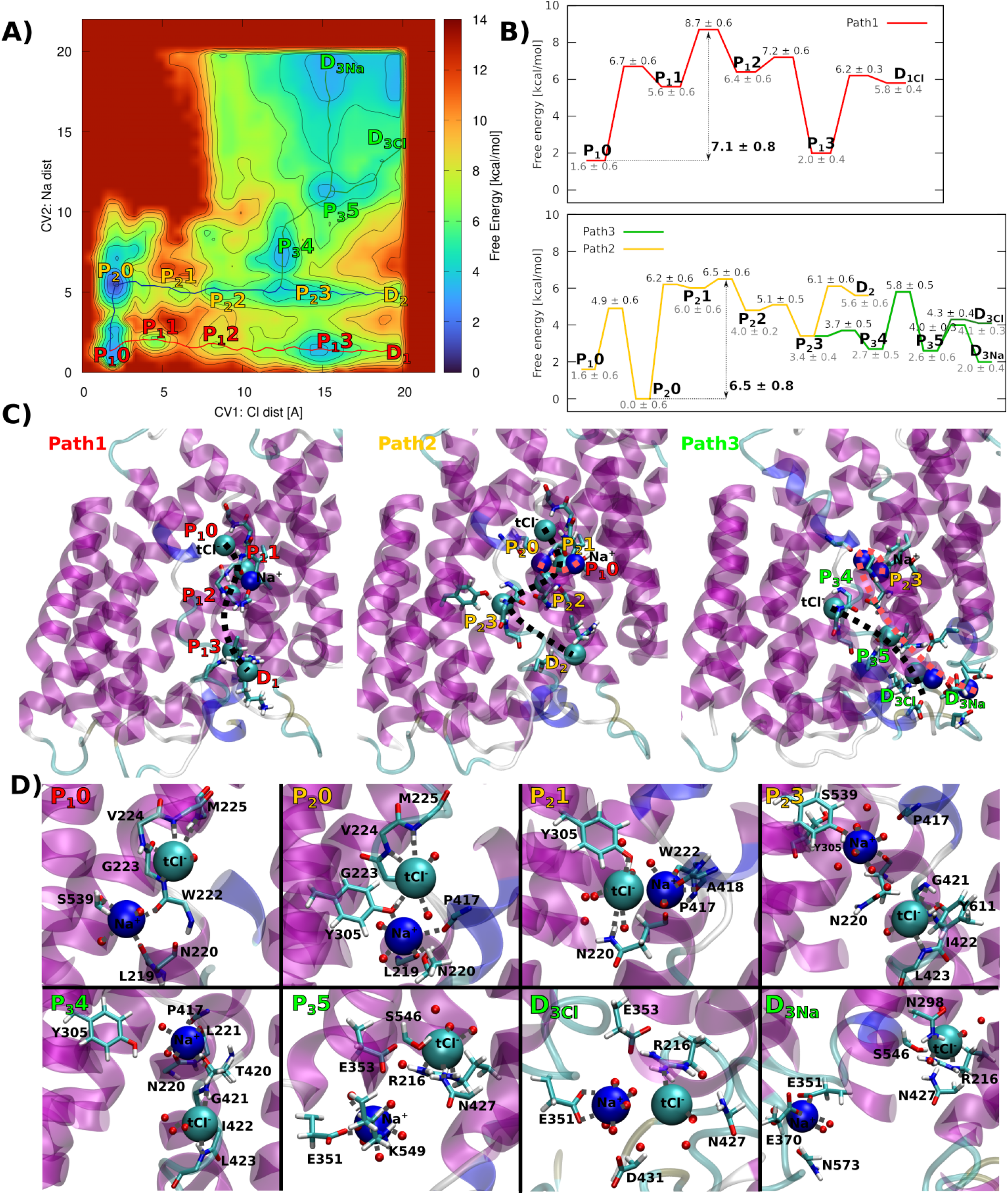
(A) Free energy surface (kcal/mol) reported as a function of the distances (Å) of the most protein buried (top, t) Cl^-^ and Na^+^ ions from their initial binding site (CV1 and CV2, respectively). The free energy isosurface is shown from blue to red color; (B) Simplified graphs showing the translocation free energy for all minima and transition states met along the Path 1 and 2/3. Free energies are reported with respect to the most stable minimum. (C) View of the tCl^-^ and Na^+^ ions dissociation pathways along the NKCC1 channel with subsequent positions of tCl^-^ and Na^+^ highlighted with black and red dashed lines, respectively. (D) Close-ups of the initial binding sites and the most relevant states visited along the most energetically viable pathway (Path 3). The coordination spheres of the Na^+^ and tClion are highlighted in gray dashed lines. The NKCC1 protein is depicted as magenta new cartoon, the Na^+^ and tCl^-^ ions as dark blue and light blue van der Waals spheres, respectively. The protein residues and water molecules forming the coordination spheres are shown in licorice and balls (without hydrogens), respectively.

#### Path 1: Translocation Mechanism of the Most Protein-Buried (top) Cl^-^ Ion

In the initial binding state (**P**_**1**_**0**, P1 stands for Path1, while the number refers to the minimum encountered along the route, Figure 5), Na^+^ is coordinated by the backbone carbonyl oxygens of Leu219, Asn220, and Trp222, and by hydroxyl of the Ser429 side-chain, while tCl^-^ is bound by the backbone amide of residues Gly223, Val224 and Met225.

**Figure 5.**
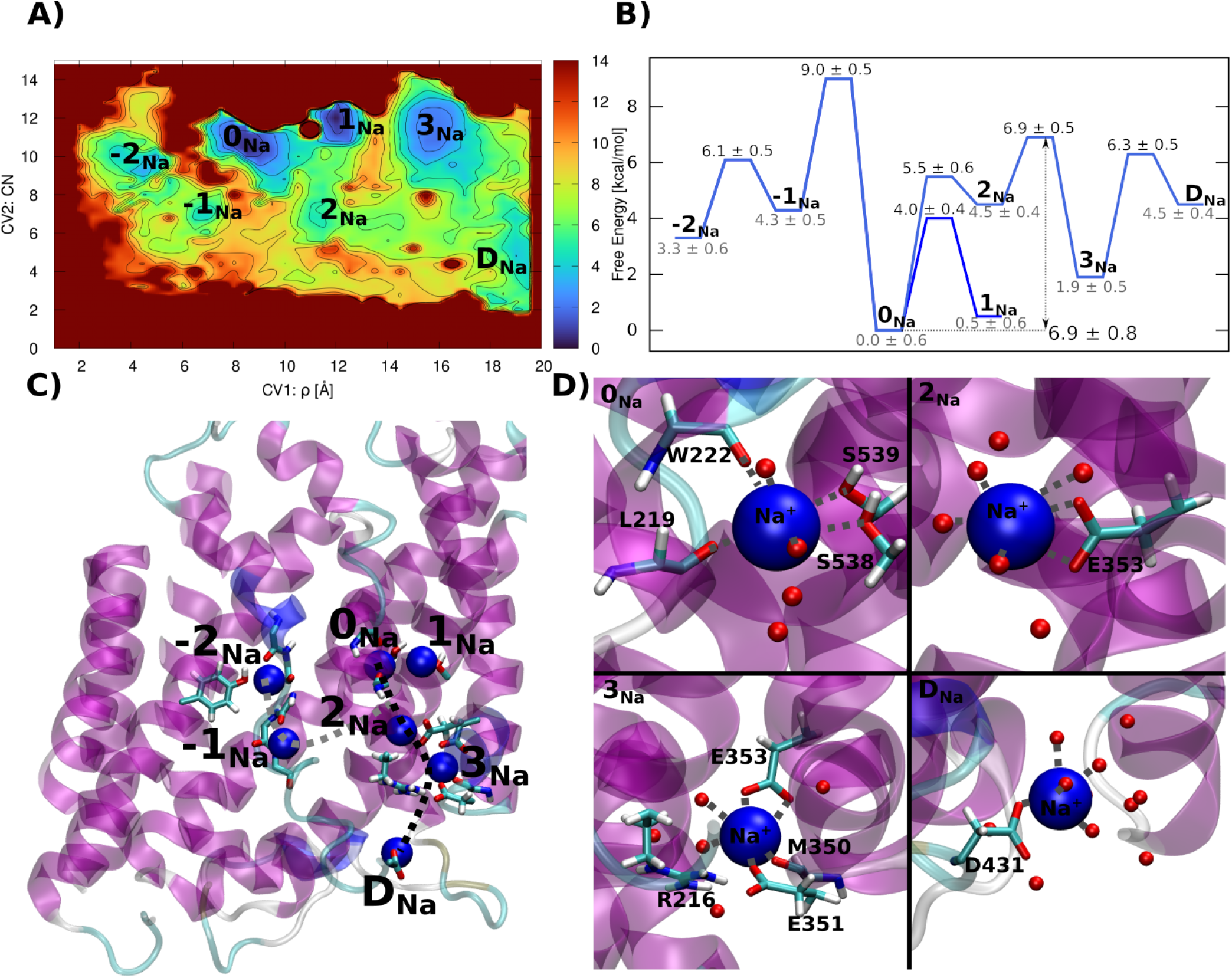
(A) Free energy surface (kcal/mol) of Na^+^ dissociation reported as a function Collective Variable (CV) 1 and CV2 defined as the distance (Å) of the ion from the protein center and of the coordination number to protein residues, respectively. The free energy isosurface is shown from blue to red color. (B) Simplified graph showing the Na^+^ dissociation free energy for all minima and transition states along the translocation path. States labeled with negative numbers refers to minima that do not enable the Na^+^ progress towards the channel exit. (C) Image showing the Na^+^ translocation path along the channel with the subsequent position of the Na^+^ ion highlighted with a black dashed line. (D) Close-up view of initial binding site and selected relevant minima visited by the Na^+^ ion during its transport. The coordination sphere of the Na^+^ ion is indicated with gray dashed lines. The NKCC1 protein is depicted as magenta new cartoons, the Na^+^ as blue van der Waals spheres, protein residues and water molecules are shown in licorice and balls (without hydrogens), respectively.

**Figure 6.**
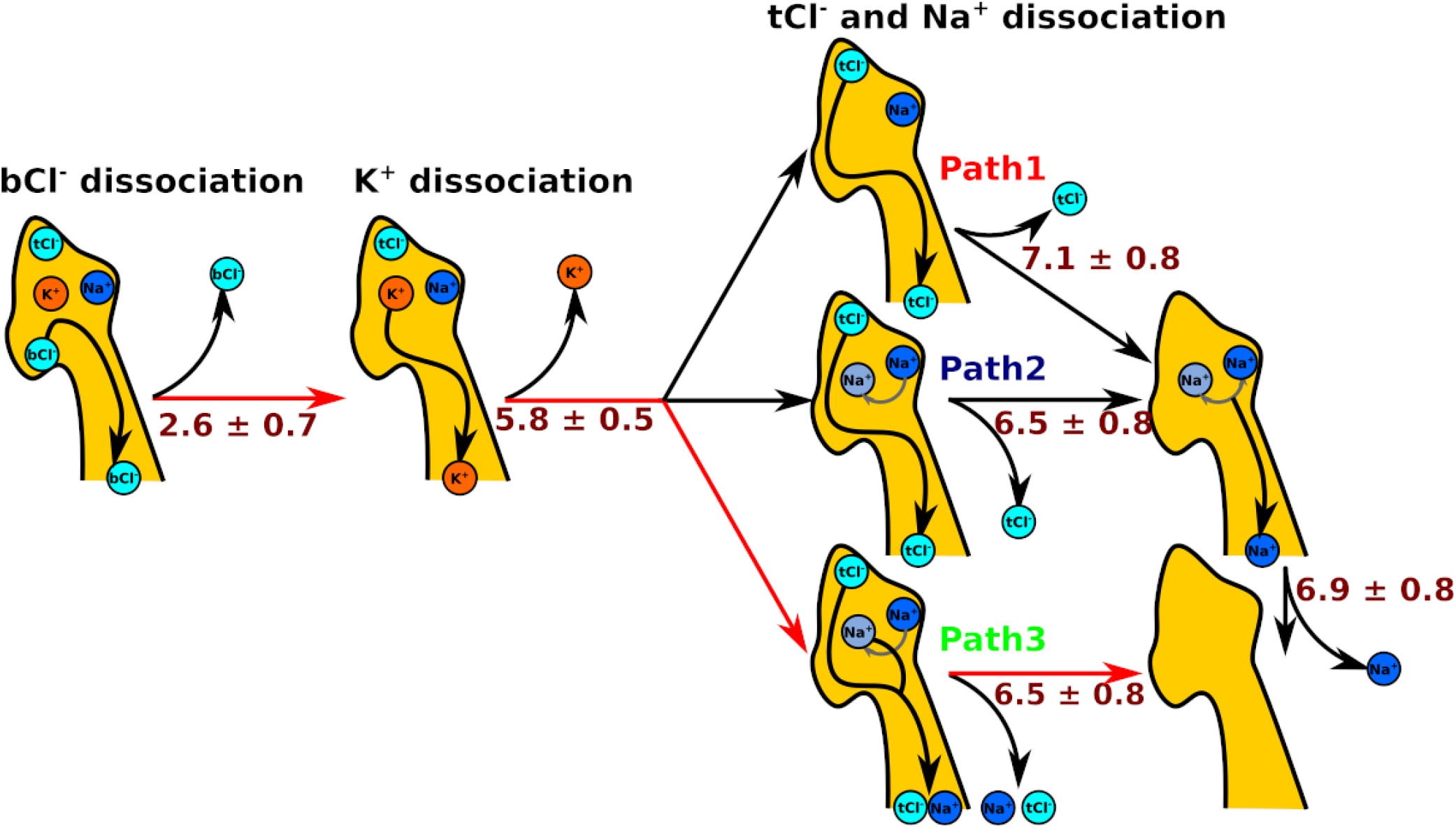
Schematic representation of the mechanism of NKCC1 ions in-cell translocation process. Ions binding cavity and the exit channel are represented as a yellow surface. The Cl^-^, Na^+^ and K^+^ ions are depicted in cyan, blue and orange spheres, with a black arrow indicating the order of dissociation events. Free energy barriers (kcal/mol) are reported in brown. The most energetically viable path is highlighted with red arrows.

While Na^+^ remains in its binding pocket, the tCl^-^ moves towards Na^+^, visiting the intermediate **P**_**1**_**1** state, through a ΔG^‡^_B1→ P11_ = 5.1 ± 0.8 kcal/mol. Here, tCl^-^ is stabilized by the electrostatic interactions with Na^+^ and is coordinated by the Tyr305 and Ser539 hydroxyl groups. Next, tCl^-^ reaches **P**_**1**_**2**, (ΔG^‡^_P11→ P12_ = 3.1 ± 0.8 kcal/mol), where it is bound only by Tyr503 hydroxyl group. By surmounting a ΔG^‡^_P12→ P13_ of 0.8 ± 0.8 kcal/mol, tCl^-^ approaches the last intermediate state **P 3**, coordinated by the amide moiety of Asn427 side-chain and the guanidinium group of Arg216. From here, tCl^-^ dissociates, through a ΔG^‡^_P13→ D1_ of 4.2 ± 0.5 kcal/mol, with the help of the Asn427 and Lys549 residues. In summary, Path 1 exclusively involves tCl^-^ dissociation with an overall ΔG^‡^_P1_ of 7.1 ± 0.8 kcal/mol and the observed metastable states (Figure S12) lie at 4.0-5 kcal/mol above the main minimum (Figure 4).

#### Path 2: Translocation Mechanism of the Most Protein-buried (top) Cl^-^ Ion Adjuvated by the Na^+^ Ion

Alternatively, Path 2 starts with Na^+^ moving to the **P**_**2**_**0** minimum (ΔG^‡^_P10→ P20_ 3.3 ± 0.6 kcal/ mol), which corresponds to the initial K^+^ binding site (**0**_**K**,_ Figure 3). This state is the main minimum of the FES (Figure 4), with Na^+^ being coordinated by the backbone carbonyl oxygen of Pro427, Asn420 and Gly421. From **P**_**2**_**0**, the tCl^-^ dissociates via three intermediate states **P**_**2**_**1-3** (Figure S12) characterized by ΔG^‡^s of 6.2 ± 0.6, 0.3 ± 0.6 and 1.1 ± 0.7 kcal/mol, respectively. The tCl^-^ in **P**_**2**_**1** and **P**_**2**_**2** binds almost identically as in the **P**_**1**_**1** and **P**_**1**_**2** sites, while the position of Na^+^ is different. Conversely, in **P**_**2**_**3** state tCl^-^ occupies the initial bCl^-^ binding site (**0**_**bCl**_, Figure 2). From **P**_**2**_**3**, three different egress routes open: (i) the tCl^-^ dissociates through **D**_**2**_ state, by overcoming a ΔG^‡^_P23→ D2_ of 2.7 ± 0.7 kcal/mol, (ii) tCl^-^ reverts to path 1, by surmounting a ΔG^‡^_P23→ P13_ = 2.4 ± 0.6 kcal/mol; (iii) or it moves to path 3, detailed in the following section. Assuming that tCl^-^ would translocate along Path 1 or 2, it will require to surmount an overall ΔG^‡^_dtcl_ of 6.5 ± 0.8 kcal/mol (Figure 4), with the mestastable states (Figure S12) lying at 3.4-4.0 kcal/mol above the main minimum **P**_**2**_**0**.

#### Path 3: Cooperative Translocation Mechanism of the top Cl^-^ and Na^+^ ions

From **P**_**2**_**3** Na^+^ moves into the **P**_**3**_**4** at negligible ΔG^‡^_P23→ P34_ of 0.3 ± 0.6 kcal/mol. This defines a third possible exit path (Path 3). In **P**_**3**_**4** Na^+^ is coordinated by the carbonyl oxygens of Leu221 and Pro417, being further stabilized by a water-mediated interaction with Tyr305 and Asn220. tCl^-^ instead remains in **P**_**2**_**3**. Next, both ions translocate to **P**_**3**_**5** (ΔG^‡^_P34→ P35_ of 3.1 ± 0.7 kcal/mol) where Na^+^ and tCl^-^ interact with the side-chains of residues Glu351 and Glu353; and Arg216, Asn427 and S546, respectively. From **P**_**3**_**5** Na^+^ moves towards **D**_**3Na**_, assisted by the side-chains of residues Glu351 and Glu575 (ΔG^‡^_P35→ D3Na_of 1.4 ± 0.7 kcal/mol), while tCl^-^ heads to **D3**_**Cl**_, aided by the Na^+^ ion and the Asn427 side-chain (ΔG^‡^_P35→ D3Cl_ of 1.7 ± 0.7 kcal/mol). Also in this case the rate-limiting step of the dissociation path is ΔG^‡^_dtCl,Na_ of 6.5 ± 0.8 kcal/mol. Within this intricate scenario it is not possible to discriminate the most energetically viable route. Therefore, we even dissected the Na^+^ translocation mechanism, assuming that this may take place after bCl^-^ dissociation along Path 2.

### Transport Mechanism of the Na^+^ Ion

After tCl^-^ dissociation, Na^+^ is not stably placed in the initial K^+^ binding site (**0**_**K**_ or **P**_**2**_**0**, Figure 4). Hence, during the MTD simulation Na^+^ visits other states in the channel interior (**-2**_**Na**_ and **-1**_**Na**_ minima, Figure 5, Figure S15), before reaching the initial Na^+^ binding site **0**_**Na**_, the global minimum of this FES. Here Na^+^ is bound by the carbonyl oxygens of residues Leu219 and Trp222 and by the hydroxyl group of Ser538 and Ser539 side-chains. Remarkably, this minimum is separated by ΔG^‡^_0→ 1_= ± 0.7 kcal/mol from a second energetically equivalent minimum **1**_**Na**_, where Na^+^ loses its coordination with carbonyl oxygens of Leu219 and Trp222. The presence of two energetically equivalent minima may provide a rationale to the difficulties of tracing the Na^+^ binding site in the cryo-EM structure.^20^ Nevertheless, the dissociation of Na^+^ proceeds from **0**_**Na**_ to **2**_**Na**_ via ΔG^‡^_0→ 2_= 5.5 ± 0.8 kcal/mol. In **2**_**Na**_ Na^+^ is exclusively coordinated by Glu353 side-chain and water molecules. Next, by surmounting a ΔG^‡^_2→ 3_= 2.4 ± 0.6 kcal/mol, Na^+^ is escorted by Glu353 to **3**_**Na**_, where it is bound by the Glu351 and Glu353 side-chains carboxylic groups, and by the carbonyl oxygen of Met350. Finally, Asp431 assists Na^+^ during its final translocation step (ΔG^‡^_3→ D_ of 4.4 ± 0.7 kcal/mol). The overall dissociation ΔG^‡^_dNa_ is 6.9 ± 0.8 kcal/mol, which is the highest of all the ions. The observed metastable states (Figure 5) lie 3-4.5 kcal/mol higher in free energy with respect to the most stable minimum **0**_**Na**_. Our results are consistent with the experimental evidence that the inward-facing conformation of NKCC1 has a higher affinity for Na^+^ then for K^+^.^18^ We also remark, that our MTD simulations, successfully employed in other studies to assess the coordination sphere of loosely bound ions,^37^ confirmed the identity of the Na^+^ binding site proposed on the basis of structural homology to other transporters.

## Discussion

According to our exhaustive characterization of all possible ions translocation paths, the most likely relative order of transport encompasses the initial in-cell delivery of bCl^-^ (ΔG^‡^_bCl_ = 2.6 ± 0.7 kcal/mol), which facilities the subsequent K^+^ egress (ΔG^‡^_K_ = 5.8 ± 0.5 kcal/mol). The remaining tCl^-^ and Na^+^ ions instead cooperatively exit the NKCC1 channel (ΔG^‡^_Na-tCl_ = 6.5 ± 0.8 kcal/mol) in the rate-limiting step of the whole in-cell ions translocation process. Nevertheless, we cannot exclude that the subsequent tCl^-^ and Na^+^ ions dissociation may also be energetically viable (ΔG^‡^ = 6.5 ± 0.8 and 6.9 ± 0.8 kcal/mol for tCl^-^ followed by Na^+^, respectively). Indeed, the calculated ΔG_d_^‡^s of both pathways are in good-agreement with experimental turnover rate of 3500 s^-1^ at 37 °C,^39,41^ corresponding to a maximum ΔG_d_^‡^ of 5 kcal/mol.

Strikingly, our translocation mechanism is supported by a wealth of biochemical and biophysical data. A recent transport model argued that NKCC1 in the outward-facing conformation first binds Cl^-^ followed by Na^+^, and then by the second Cl^-^ and K^+^ ions.^18^ This experimental study also demonstrated that the NKCC1 in the inward-facing conformation has lower binding affinities for the K^+^ and one Cl^-^ ions as compared to that of Na^+^ and the second Cl^-^ ions. Nevertheless, while the experimental evidences clearly indicate that K^+^ leaves before Na^+^,^18^ the relative order of dissociation between positively and negatively charged ions was unclear.

Consistently with these data, our simulations demonstrate that K^+^ precedes Na^+^ translocation. Additionally, we unambiguously disclose that bCl^-^, contributing to properly shaping the K^+^ and Na^+^ binding sites, must dissociate as the first ion. Our mechanism is consistent with the experimental evidences of a coupled translocation of positively and negatively charged ions as a necessary requirement to achieve an electroneutral transport.

Our translocation routes and mechanism also contribute to assess the role of several NKCC1 residues (reported in Table I) pinpointed in mutagenesis studies. A decreased transport activity was experimentally observed by mutating: (i) Tyr611 and Tyr454, which in our simulations, are involved in bCl^-^ translocation; (ii) Asn220, which stabilizes bCl^-^ and mediates the binding of K^+^; (iii) Asn427, which takes part in the final translocation steps of bCl^-^, K^+^ and tCl^-^;^21^ (iv) Arg216, which engages a salt-bridge with Glu353. This latter residue lines the dissociation pathways of all ions.38 Finally, several amino acids lying along the helix (H)3 were indicated to affect NKCC1 function. ^39^ Among these Tyr305 forms the K^+^ and tCl^-^ binding sites, while Asn298 coordinates the K^+^ and bCl^-^ ions and assists the initial step of the bCl^-^ translocation. Finally, Asn298 mutation was experimentally demonstrated to decrease the affinity of all ions, consistently with its role as a plug for the binding and stability of all ions observed in our simulations. These data, besides compellingly supporting the reliability of our findings, also supply a meticulous atomic-level mapping for future mutagenesis studies.

**Table 1.** Key NKCC1 residues identified in our ion translocation paths and/or in previous biophysical experimental studies. Only residues pertaining to ions translocation from the NKCC1 inward-facing conformation are listed.

| NKCC1 residue | Role | Experimental confirmation |
| --- | --- | --- |
| R216 | Intracellular gate; salt bridge with E353; coordinates $\text{Cl}^-$ | R216A; reduced transport; ref.<sup>38</sup> |
| N220 | Coordinates bCl<sup>-</sup>, K<sup>+</sup>, tCl<sup>-</sup>, Na<sup>+</sup> | N220A; diminished activity; ref.<sup>20</sup> |
| F294 | Channel lining; contact with K<sup>+</sup> | F294W; reduced ion affinities; ref.<sup>39</sup> |
| A297 | Stability; indirect effect via N298 | A297W; reduced transport activity; ref.<sup>39</sup> |
| N298 | Coordinates K<sup>+</sup>; water mediated interaction with Na<sup>+</sup>, Cl<sup>-</sup> | N298W; loss of function; ref.<sup>39</sup> |
| Y305 | Coordinates K<sup>+</sup>, tCl<sup>-</sup>; binding site | Y383F, Y383C, Y383W; reduced transport; ref.<sup>39</sup> |
| E351 | Channel exit; coordinates Na<sup>+</sup> | - |
| E353 | Intracellular gate; salt bridge with R216; coordinates K<sup>+</sup>, Na<sup>+</sup> | - |
| P417 | Helix kink; K<sup>+</sup> binding site | P417C; loss of function; ref.<sup>21,40</sup> |
| A418 | Helix kink; K<sup>+</sup> binding site | A418C; loss of function; ref.<sup>21,40</sup> |
| T420 | Conformational switch; coordinates K<sup>+</sup>, Na<sup>+</sup> | T420C; decreased transport activity; ref.<sup>21</sup> |
| N427 | Coordinates K<sup>+</sup>, bCl<sup>-</sup>, tCl<sup>-</sup> | N427C; ref.<sup>21</sup> |
| Y454 | Coordinates tCl<sup>-</sup>; part of tCl<sup>-</sup> site | Y454A diminished activity; ref.<sup>20</sup> |
| K549 | Channel exit; interacts with Cl<sup>-</sup> | K549A, C, D; loss of function; ref.<sup>21</sup> |
| Y611 | Coordinates Cl<sup>-</sup>; part of bCl<sup>-</sup> site | Y611A; diminished activity; ref.<sup>20</sup> |

What is more, a volume cavity analysis of the human NKCC1 transmembrane domain cryoEM structure identified three possible extracellular-entrance and three intracellular-exit channels.^38^ Building on this observation it was speculated that different ions may travel along distinct binding/ dissociation routes. Conversely, our simulations unprecedentedly reveal that all ions are transported by NKCC1 via the same channel (Figure S16) labeled as IV in ref.^38^ In this context, it is tempting to argue that the residues Asn220, Asn298 and Asn427, common to the different ion translocation pathways, and demonstrated by mutagenesis studies to play key roles in ions transport, could act as promiscuous residues able to mediate the translocation of both positively and negatively charged ions owing to their ambivalent side-chains. In compelling support of this argument these three asparagine residues are conserved in NKCC2, while Asn220 and Asn427 are conserved in KCC1-4 channels as well (Figure S17).

In summary, this study represents the first attempt of tracing the atomic-level details, and the kinetic and thermodynamic signature of NKCC1 ions transport mechanism. Consistently with a wealth of biophysical and biochemical data our outcomes strikingly unlock the relative order of ions translocation, and elucidate the molecular terms of their cooperativity.^18^ Ions transport occurs through a rugged free energy surface hallmarked by the presence of multiple metastable states connected by multi-branched pathways. The rate-limiting step (ΔG^‡^ = 6.5 ± 0.8 kcal/mol) of the whole transport process, underlying the cooperative dissociation of Na^+^ and tCl^-^ ions, is consistent with the experimental turnover-rate.^41^ Stunningly, all ions reach the cytosol by traversing the same exit channel, likely owing to the side-chain promiscuity of several asparagine residues, conserved in the CCC family, that aptly adapt to bind either positively or negatively charged ions. Owing to the overwhelming implications of NKCC1-mediated Cl^-^ dishomeostasis in a variety of human diseases, ranging to neuropathies to renal disturbances and cancer, the atomic-level dissection of the NKCC1 transport mechanism supplied here advances our understanding of the whole CCC family function and sets a conceptual basis for future drug-design efforts aiming at devising selective NKCC1-targeting inhibitors.

## Acknowledgments

This work has been conducted with the financial support of the project ‘Against bRain cancEr: finding personalized therapies with in Silico and in vitro strategies‘ (ARES) CUP: D93D19000020007 POR FESR 2014 2020 - 1.3.b -Friuli Venezia Giulia. AM thanks the financial support of the Italian Association for Cancer Research (IG grant 24514).

## Notes

### Competing Interest Statement

The authors have declared no competing interest.

## References

(1) Russell, J. M. Sodium-Potassium-Chloride Cotransport. Physiological reviews 2000, 80 (1), 211–276.

(2) Kaila, K.; Price, T. J.; Payne, J. A.; Puskarjov, M.; Voipio, J. Cation-Chloride Cotransporters in Neuronal Development, Plasticity and Disease. Nat Rev Neurosci 2014, 15 (10), 637–654. https://doi.org/10.1038/nrn3819.

(3) Markadieu, N.; Delpire, E. Physiology and Pathophysiology of SLC12A1/2 Transporters. Pflugers Arch - Eur J Physiol 2014, 466 (1), 91–105. https://doi.org/10.1007/s00424-013-1370-5.

(4) Lytle, C.; Forbush 3rd, B. Regulatory Phosphorylation of the Secretory Na-K-Cl Cotransporter: Modulation by Cytoplasmic Cl. American Journal of Physiology-Cell Physiology 1996, 270 (2), C437–C448.

(5) Garneau, A. P.; Slimani, S.; Fiola, M. J.; Tremblay, L. E.; Isenring, P. Multiple Facets and Roles of Na+ -K+ -Cl− Cotransport: Mechanisms and Therapeutic Implications. Physiology 2020, 35 (6), 415–429. https://doi.org/10.1152/physiol.00012.2020.

(6) Schulte, J. T.; Wierenga, C. J.; Bruining, H. Chloride Transporters and GABA Polarity in Developmental, Neurological and Psychiatric Conditions. Neuroscience & Biobehavioral Reviews 2018, 90, 260–271. https://doi.org/10.1016/j.neubiorev.2018.05.001.

(7) Contestabile, A.; Magara, S.; Cancedda, L. The GABAergic Hypothesis for Cognitive Disabilities in down Syndrome. Frontiers in cellular Neuroscience 2017, 11, 54.

(8) Cellot, G.; Cherubini, E. GABAergic Signaling as Therapeutic Target for Autism Spectrum Disorders. Frontiers in pediatrics 2014, 2, 70.

(9) Gamba, G. Molecular Physiology and Pathophysiology of Electroneutral Cation-Chloride Cotransporters. Physiological Reviews 2005, 85 (2), 423–493. https://doi.org/10.1152/physrev.00011.2004.

(10) Gagnon, K. B.; Delpire, E. Physiology of SLC12 Transporters: Lessons from Inherited Human Genetic Mutations and Genetically Engineered Mouse Knockouts. American Journal of Physiology-Cell Physiology 2013, 304 (8), C693–C714. https://doi.org/10.1152/ajpcell.00350.2012.

(11) Luo, L.; Wang, J.; Ding, D.; Hasan, M. N.; Yang, S.-S.; Lin, S.-H.; Schreppel, P.; Sun, B.; Yin, Y.; Erker, T.; Sun, D. Role of NKCC1 Activity in Glioma K+ Homeostasis and Cell Growth: New Insights With the Bumetanide-Derivative STS66. Front. Physiol. 2020, 11, 911. https://doi.org/10.3389/fphys.2020.00911.

(12) Schrier, R. W. Use of Diuretics in Heart Failure and Cirrhosis. In Seminars in nephrology; Elsevier, 2011; Vol. 31, pp 503–512.

(13) Edwards, D. A.; Shah, H. P.; Cao, W.; Gravenstein, N.; Seubert, C. N.; Martynyuk, A. E. Bumetanide Alleviates Epileptogenic and Neurotoxic Effects of Sevoflurane in Neonatal Rat Brain. Anesthesiology: The Journal of the American Society of Anesthesiologists 2010, 112 (3), 567–575.

(14) Auer, T.; Schreppel, P.; Erker, T.; Schwarzer, C. Functional Characterization of Novel Bumetanide Derivatives for Epilepsy Treatment. Neuropharmacology 2020, 162, 107754. https://doi.org/10.1016/j.neuropharm.2019.107754.

(15) Damier, P.; Hammond, C.; Ben-Ari, Y. Bumetanide to Treat Parkinson Disease: A Report of 4 Cases. Clinical neuropharmacology 2016, 39 (1), 57–59.

(16) Bisha, I.; Magistrato, A. The Molecular Mechanism of Secondary Sodium Symporters Elucidated through the Lens of the Computational Microscope. RSC Adv. 2016, 6 (12), 9522–9540. https://doi.org/10.1039/C5RA22131E.

(17) Haas, M.; Forbush III, B. The Na-K-Cl Cotransporter of Secretory Epithelia. Annual review of physiology 2000, 62 (1), 515–534.

(18) Delpire, E.; Gagnon, K. B. Kinetics of Hyperosmotically Stimulated Na-K-2Cl Cotransporter in Xenopus Laevis Oocytes. American Journal of Physiology-Cell Physiology 2011, 301 (5), C1074–C1085.

(19) Gagnon, K. B.; Delpire, E. Molecular Determinants of Hyperosmotically Activated NKCC1-Mediated K+/K + Exchange: Volume-Sensitive NKCC1-Mediated K+/K+ Exchange. The Journal of Physiology 2010, 588 (18), 3385– 3396. https://doi.org/10.1113/jphysiol.2010.191932.

(20) Chew, T. A.; Orlando, B. J.; Zhang, J.; Latorraca, N. R.; Wang, A.; Hollingsworth, S. A.; Chen, D.-H.; Dror, R. O.; Liao, M.; Feng, L. Structure and Mechanism of the Cation–Chloride Cotransporter NKCC1. Nature 2019, 572 (7770), 488–492. https://doi.org/10.1038/s41586-019-1438-2.

(21) Zhang, S.; Zhou, J.; Zhang, Y.; Liu, T.; Friedel, P.; Zhuo, W.; Somasekharan, S.; Roy, K.; Zhang, L.; Liu, Y.; Meng, X.; Deng, H.; Zeng, W.; Li, G.; Forbush, B.; Yang, M. The Structural Basis of Function and Regulation of Neuronal Cotransporters NKCC1 and KCC2. Commun Biol 2021, 4 (1), 226. https://doi.org/10.1038/s42003-021-01750-w.

(22) Webb, B.; Sali, A. Comparative Protein Structure Modeling Using MODELLER. Current protocols in bioinformatics 2016, 54 (1), 5–6.

(23) Schrödinger Release 2013-3: Maestro, Version 9.6; Schrödinger, LLC: New York, NY, 2013.

(24) Jo, S.; Kim, T.; Iyer, V. G.; Im, W. CHARMM-GUI: A Web-Based Graphical User Interface for CHARMM. Journal of computational chemistry 2008, 29 (11), 1859–1865.

(25) Wu, E. L.; Cheng, X.; Jo, S.; Rui, H.; Song, K. C.; Dávila-Contreras, E. M.; Qi, Y.; Lee, J.; Monje-Galvan, V.; Venable, R. M.; others. CHARMM-GUI Membrane Builder toward Realistic Biological Membrane Simulations. Journal of computational chemistry 2014, 35 (27), 1997–2004.

(26) Maier, J. A.; Martinez, C.; Kasavajhala, K.; Wickstrom, L.; Hauser, K. E.; Simmerling, C. Ff14SB: Improving the Accuracy of Protein Side Chain and Backbone Parameters from Ff99SB. Journal of Chemical Theory and Computation 2015, 11 (8), 3696–3713. https://doi.org/10.1021/acs.jctc.5b00255.

(27) Dickson, C. J.; Madej, B. D.; Skjevik, Å.A.; Betz, R. M.; Teigen, K.; Gould, I. R.; Walker, R. C. Lipid14: The Amber Lipid Force Field. J. Chem. Theory Comput. 2014, 10 (2), 865–879. https://doi.org/10.1021/ct4010307.

(28) Jorgensen, W. L.; Chandrasekhar, J.; Madura, J. D.; Impey, R. W.; Klein, M. L. Comparison of Simple Potential Functions for Simulating Liquid Water. The Journal of Chemical Physics 1983, 79 (2), 926. https://doi.org/10.1063/1.445869.

(29) Joung, I. S.; Cheatham, T. E. Determination of Alkali and Halide Monovalent Ion Parameters for Use in Explicitly Solvated Biomolecular Simulations. The Journal of Physical Chemistry B 2008, 112 (30), 9020–9041. https://doi.org/10.1021/jp8001614.

(30) Abraham, M. J.; Murtola, T.; Schulz, R.; Páll, S.; Smith, J. C.; Hess, B.; Lindahl, E. GROMACS: High Performance Molecular Simulations through Multi-Level Parallelism from Laptops to Supercomputers. SoftwareX 2015, 1, 19–25.

(31) Lindahl; Abraham; Hess Spoel, V.D. GROMACS 2020.4 Manual. 2020. https://doi.org/10.5281/ZENODO.4054996.

(32) Tribello, G. A.; Bonomi, M.; Branduardi, D.; Camilloni, C.; Bussi, G. PLUMED 2: New Feathers for an Old Bird. Computer Physics Communications 2014, 185 (2), 604–613. https://doi.org/10.1016/j.cpc.2013.09.018.

(33) Bonomi, M.; Bussi, G.; Camilloni, C.; Tribello, G. A.; Banáš, P.; Barducci, A.; Bernetti, M.; Bolhuis, P. G.; Bottaro, S.; Branduardi, D.; others. Promoting Transparency and Reproducibility in Enhanced Molecular Simulations. Nature methods 2019, 16 (8), 670–673.

(34) Balusek, C.; Hwang, H.; Lau, C. H.; Lundquist, K.; Hazel, A.; Pavlova, A.; Lynch, D. L.; Reggio, P. H.; Wang, Y.; Gumbart, J. C. Accelerating Membrane Simulations with Hydrogen Mass Repartitioning. J. Chem. Theory Comput. 2019, 15 (8), 4673–4686. https://doi.org/10.1021/acs.jctc.9b00160.

(35) Capelli, R.; Carloni, P.; Parrinello, M. Exhaustive Search of Ligand Binding Pathways via Volume-Based Metadynamics. J. Phys. Chem. Lett. 2019, 10 (12), 3495–3499. https://doi.org/10.1021/acs.jpclett.9b01183.

(36) Dutzler, R.; Campbell, E. B.; MacKinnon, R. Gating the Selectivity Filter in ClC Chloride Channels. Science 2003, 300 (5616), 108–112.

(37) Bisha, I.; Laio, A.; Magistrato, A.; Giorgetti, A.; Sgrignani, J. A Candidate Ion-Retaining State in the Inward-Facing Conformation of Sodium/Galactose Symporter: Clues from Atomistic Simulations. J. Chem. Theory Comput. 2013, 9 (2), 1240–1246. https://doi.org/10.1021/ct3008233.

(38) Yang, X.; Wang, Q.; Cao, E. Structure of the Human Cation–Chloride Cotransporter NKCC1 Determined by Single-Particle Electron Cryo-Microscopy. Nat Commun 2020, 11 (1), 1016. https://doi.org/10.1038/s41467-020-14790-3.

(39) Somasekharan, S.; Tanis, J.; Forbush, B. Loop Diuretic and Ion-Binding Residues Revealed by Scanning Mutagenesis of Transmembrane Helix 3 (TM3) of Na-K-Cl Cotransporter (NKCC1). J. Biol. Chem. 2012, 287 (21), 17308–17317. https://doi.org/10.1074/jbc.M112.356014.

(40) Dehaye, J.-P.; Nagy, A.; Premkumar, A.; Turner, R. J. Identification of a Functionally Important Conformation-Sensitive Region of the Secretory Na+-K+-2Cl-Cotransporter (NKCC1). Journal of Biological Chemistry 2003, 278 (14), 11811–11817.

(41) Haas, M.; Forbush, B. [3H]Bumetanide Binding to Duck Red Cells. Correlation with Inhibition of (Na + K + 2Cl) Co-Transport. Journal of Biological Chemistry 1986, 261 (18), 8434–8441. https://doi.org/10.1016/S0021-9258(19)83931-5., 3495–3499 (2019).

