## Supplementary Information for "All-atom Simulations Uncover the Molecular Terms of NKCC1 Transport Mechanism"

CNR-IOM c/o International School for Advanced studies (SISSA/ISAS), via Bonomea 265, 34136, Trieste, Italy

### Supplementary Results

#### *Molecular dynamics simulations of complete dimeric NKCC1 model*

We performed 500 ns of Molecular Dynamics (MD) simulations of a complete NKCC1 model containing a dimeric transmembrane domain and the C-term domain and of a reduced model comprising a momoner of transmembrane domain, as used also in ref. (1) (Figure S1). A comparison of root mean square deviation (RMSD) of the backbone atoms of the transmembrane helices of the complete and reduced models evidences that NKCC1 maintains the structure of its transmembrane domain even in the reduced model system. As well, the orientation of the monomeric transmembrane domain remains the same of larger dimeric model (Figure S2) as shown by the value of the angle between the transmembrane helix, helix 6, and axis perpendicular to the membrane plain. Building on these results and to reduce the computational cost of the simulations, we exploited the reduced model system to unravel the molecular mechanism of the ions transport. To further inspect the structural stability of the reduced model we also performed 500 ns of MD simulations in the presence of 150 mM KCl ionic strength in place of NaCl. The different salt did not affect the structure of the NKCC1 channel.

#### Structural stability of the ions binding sites

After having validated the structural stability of the reduced $NKCC1_{cut}$ model as compared to the larger one (Figures S1-S3), we inspected the stability of each ion binding site (Figures S4-S6) by performing exhaustive MD simulations in the present of different ions. In the MD simulations of the fully ions-loaded $NKCC1_{cut}$ model all ions preserved most of coordination ligands proposed on the basis of the cryo-EM structure, with water molecules completing their hexahedral coordination sphere. Specifically, the most cytosol-exposed $Cl^-$ ion (here after named as bottom $Cl^-$, $bCl^-$, Figure S4A) was coordinated by backbone amide hydrogens of residues Gly421, Leu522 and Leu423; and the hydroxyl hydrogen of Tyr611; the $K^+$ ion (Figure S5A) was bound by the backbone carbonyl oxygens of residues Ile221, Pro417, Ala418 and Thr420. The coordination with carbonyl oxygen of Asn220 and hydroxyl oxygens of Tyr305 and Thr420, suggested in the cryo-EM structure, were instead unstable with the O-$K^+$ distances fluctuating between 2.6 and 4.0 Å during the MD simulations.

The $Na^+$ ion, placed in the binding site proposed on the basis of structural homology to other transporters (1), remained coordinated by the carbonyl oxygens of residues Leu219 and Trp222, and by the Ser539 side-chain (Figure S6A). However, the experimentally proposed coordination with backbone and side-chain oxygens of Ala535 and Ser538, respectively, was not retained during MD simulation. The second inner-most $Cl^-$ ion (hereafter named as top $Cl^-$, $tCl^-$), was coordinated by backbone amide hydrogens of residues Gly223, Val224 and Met225 and by the Tyr454 hydroxyl hydrogen (Figure S4B). Our model, therefore, reliably reproduces the experimentally proposed ion binding sites. All ions are hexacoordinated through-out our simulations with water molecules completing the coordinate sphere in the absence of suitable protein residues.

The NKCC1 structure is affected by the presence of different ions (Figure S3). Indeed, the Root Mean Square Deviation (RMSD) of the ions' binding sites (as defined in Table S1) discloses that while the two $Cl^-$ ions binding sites are stable regardless of the number and types of ions (Figure S4) bound to NKCC1, the $K^+$ binding site (Figure S5), linked to the $bCl^-$ binding site via the G421 residue, becomes more flexible (less stable) in the absence of $bCl^-$ (Figure S5). As well, the $Na^+$ binding site fully forms only in the presence of all ions (Figure S6).

### Supplementary Figures

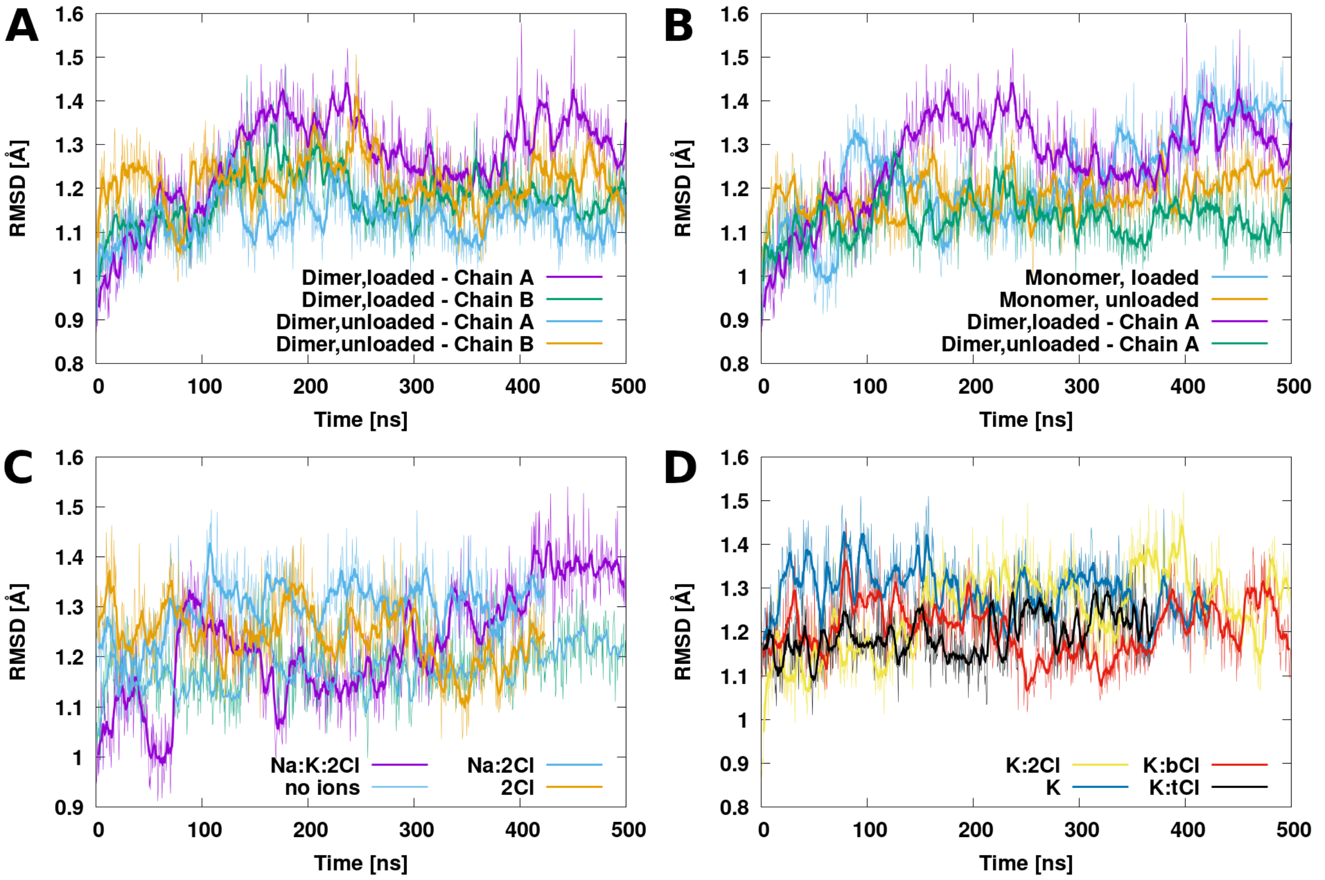

**Figure S1.** Root Mean Square Deviation (RMSD) (Å) vs simulation time (ns) for the transmembrane part of the dimeric and reduced NKCC1 model system in the presence of different ions. The two transmembrane domains of the dimeric model are labeled as chain A and B, respectively. A) Comparison of ions-loaded and empty transmembrane domains of the dimeric model. B) Comparison of loaded and unloaded transmembrane domain of the reduced monomeric model and dimeric model. C) Comparison of reduced monomeric model with different bound ions: Violet line refers to the fully loaded $Na^+$:$K^+$:2$Cl^-$ model, cyan line to the $Na^+$:2$Cl^-$-loaded model, orange line to the two $Cl^-$-loaded model and green line corresponds to the apo NKCC1 model. D) Comparison of the monomeric model with different bound ions: Yellow line refers to the $K^+$:2$Cl^-$-loaded model, blue line to the $K^+$-loaded model, red line to the $K^+$: bottom(b) $Cl^-$-loaded model and black to the $K^+$:top(t)$Cl^-$-loaded model.

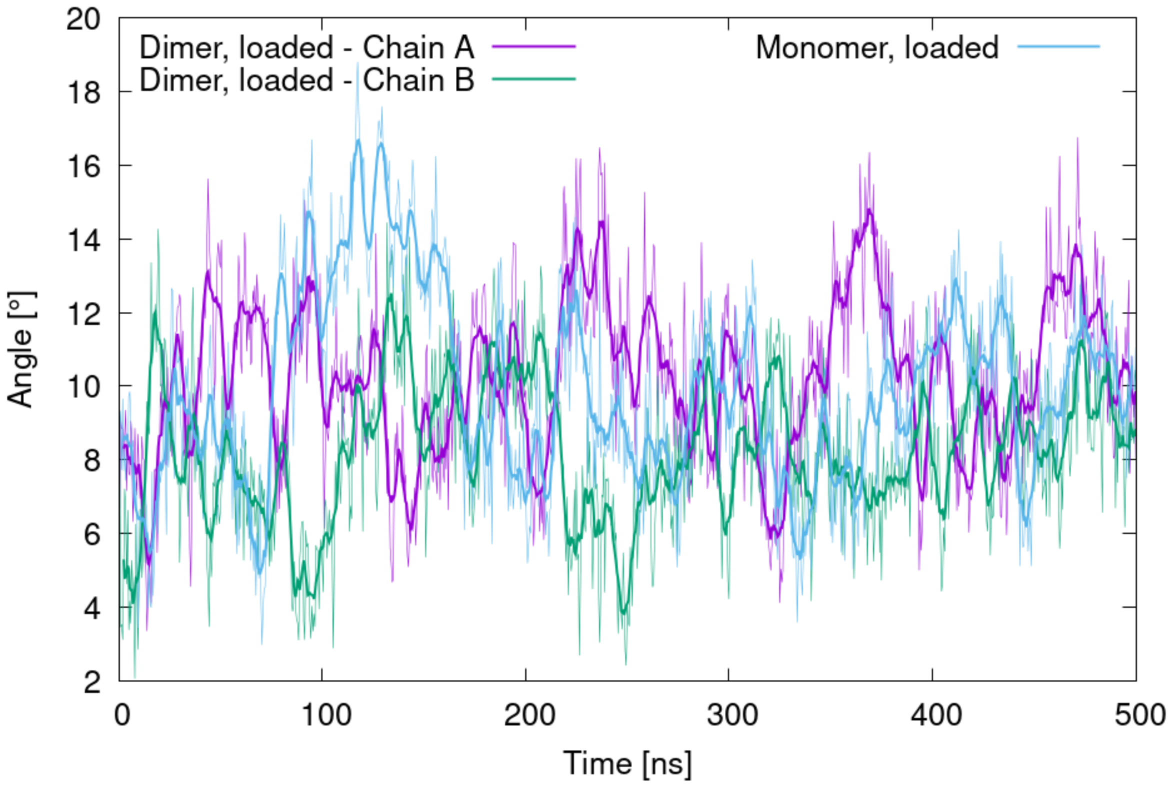

**Figure S2.** Angle (deg) between the helix 6 (defined by vector between residues 437 and 463) and the membrane plane (defined as the z-vector of the simulation cell) in the fully loaded dimeric model (purple and green lines for chain A and B, respectively) and fully loaded reduced monomeric model system (cyan line) of NKCC1. The helix is at the core of the NKCC1 transmembrane domain and it is therefore exploited as to assess the protein orientation with respect to the membrane axis.

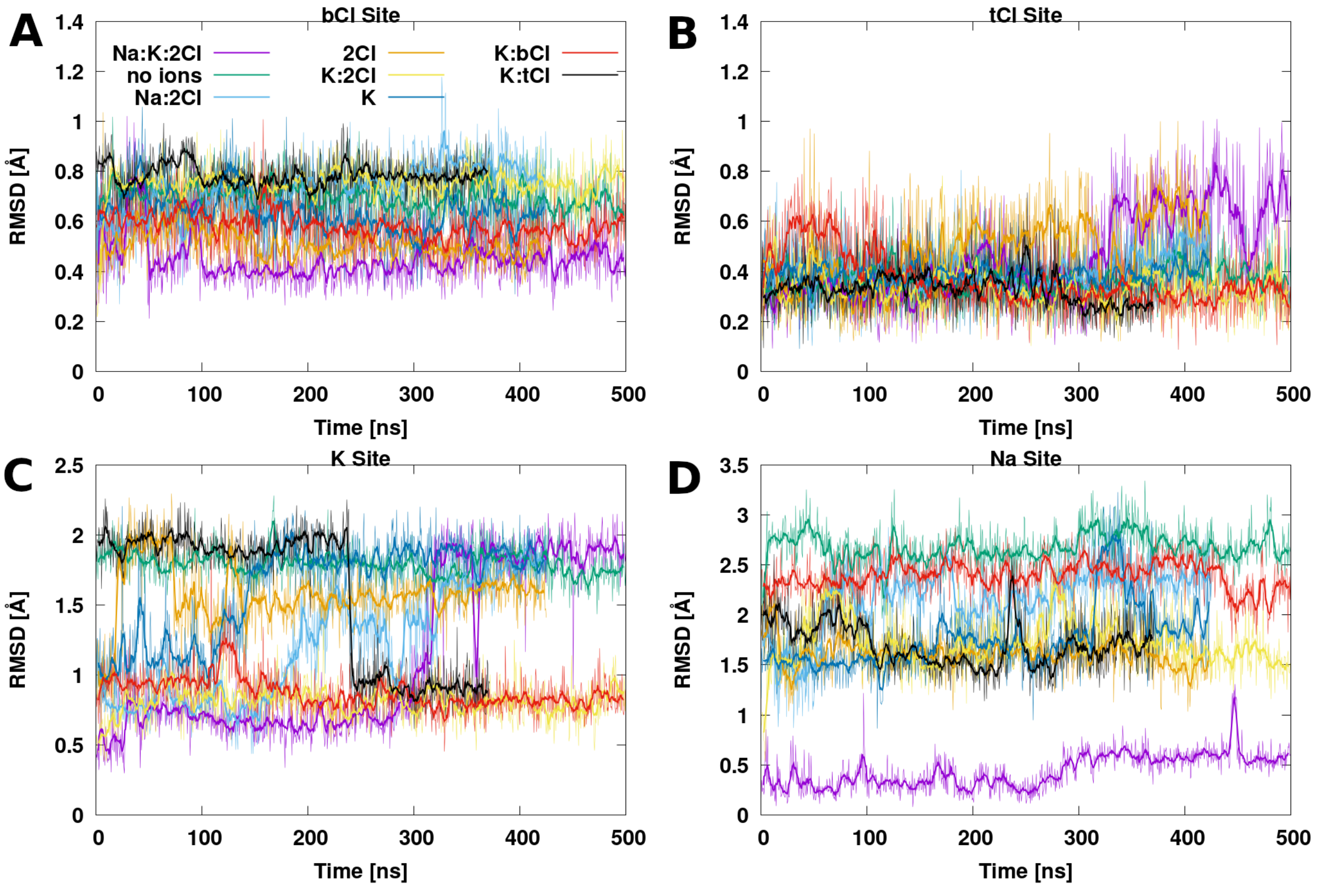

**Figure S3.** Root Mean Square Deviation (RMSD) (Å) vs simulation time (ns) of the ion binding site residues (as defined in Table S1) calculated from molecular dynamics simulation trajectory of NKCC1 reduced monomeric model with different ion content. Violet line refers to the fully loaded $Na^+$:$K^+$:2$Cl^-$ model, cyan line to the $Na^+$:2$Cl^-$-loaded model, orange line to the two $Cl^-$-loaded model, yellow line to the $K^+$:2$Cl^-$-loaded model, blue line to the $K^+$-loaded model, red line to the $K^+$:bottom(b)$Cl^-$-loaded model, black line to the $K^+$:top(t)$Cl^-$-loaded model and green line corresponds to the apo NKCC1 model.

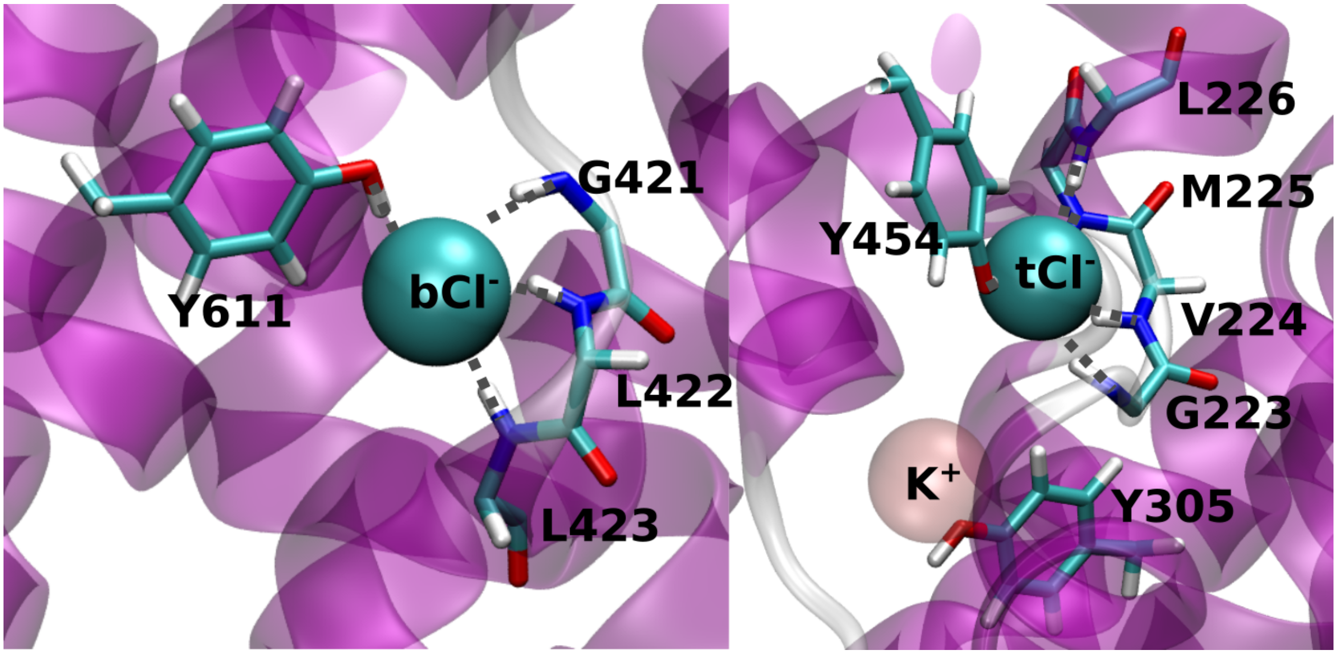

**Figure S4.** Close-up view of bottom (b)$Cl^-$ and top (t)$Cl^-$ initial binding sites. The NKCC1 protein is depicted as magenta new cartoons, the b$Cl^-$ and $K^+$ are depicted ions as cyan and pink van deer Waals spheres. Residues forming the ions coordination sphere are are shown in licorice. Water molecules completing the hexahedral coordination sphere are not shown for clarity.

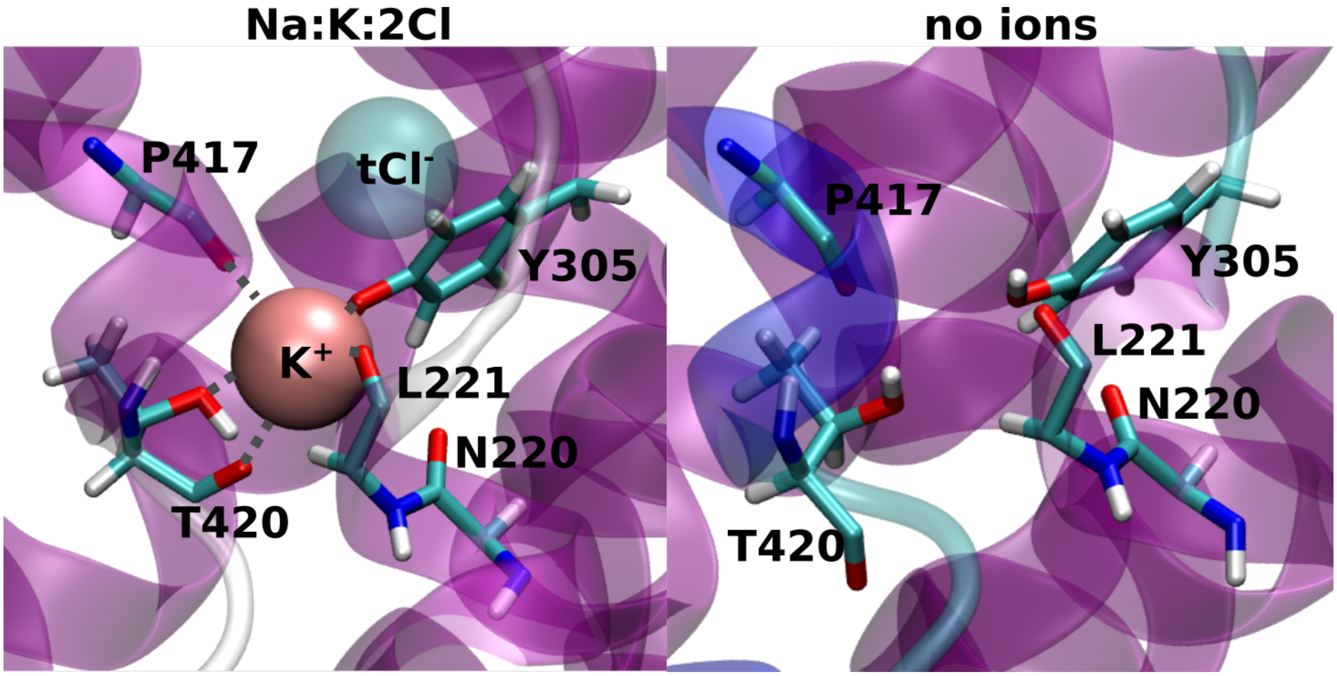

**Figure S5.** Close-up view of $K^+$ initial binding site in the presence (left panel) and in the absence of the $K^+$ and top (t)$Cl^-$ ions (right panel). The NKCC1 protein is depicted as magenta new cartoons, the t$Cl^-$ and $K^+$ ions are represented as cyan and pink van der Waals spheres. Residues defining the ions coordination sphere are shown in licorice. Water molecule completing the hexahedral coordination sphere are omitted for clarity reasons.

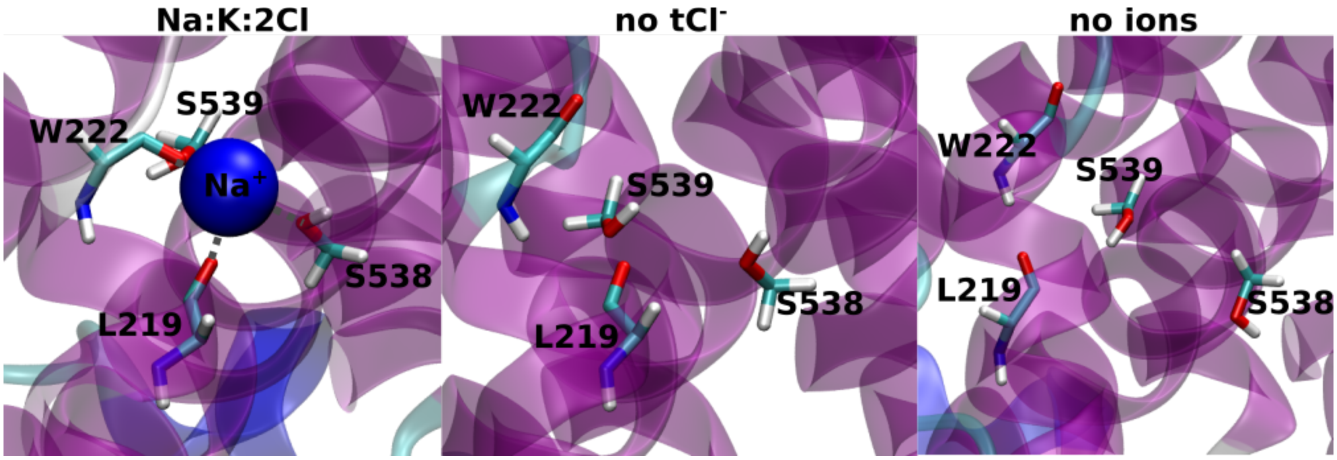

**Figure S6**. Close-up views of $Na^+$ binding site in the presence of all ions (left panel) and absence of the top (t)$Cl^-$ ion (central panel) and in the absence of all ions (right panel). The NKCC1 protein is depicted as magenta new cartoons, the $Na^+$ ion is shown as blue sphere, relevant residues are depicted in licorice. Water molecule completing the hexahedral coordination sphere are omitted for clarity.

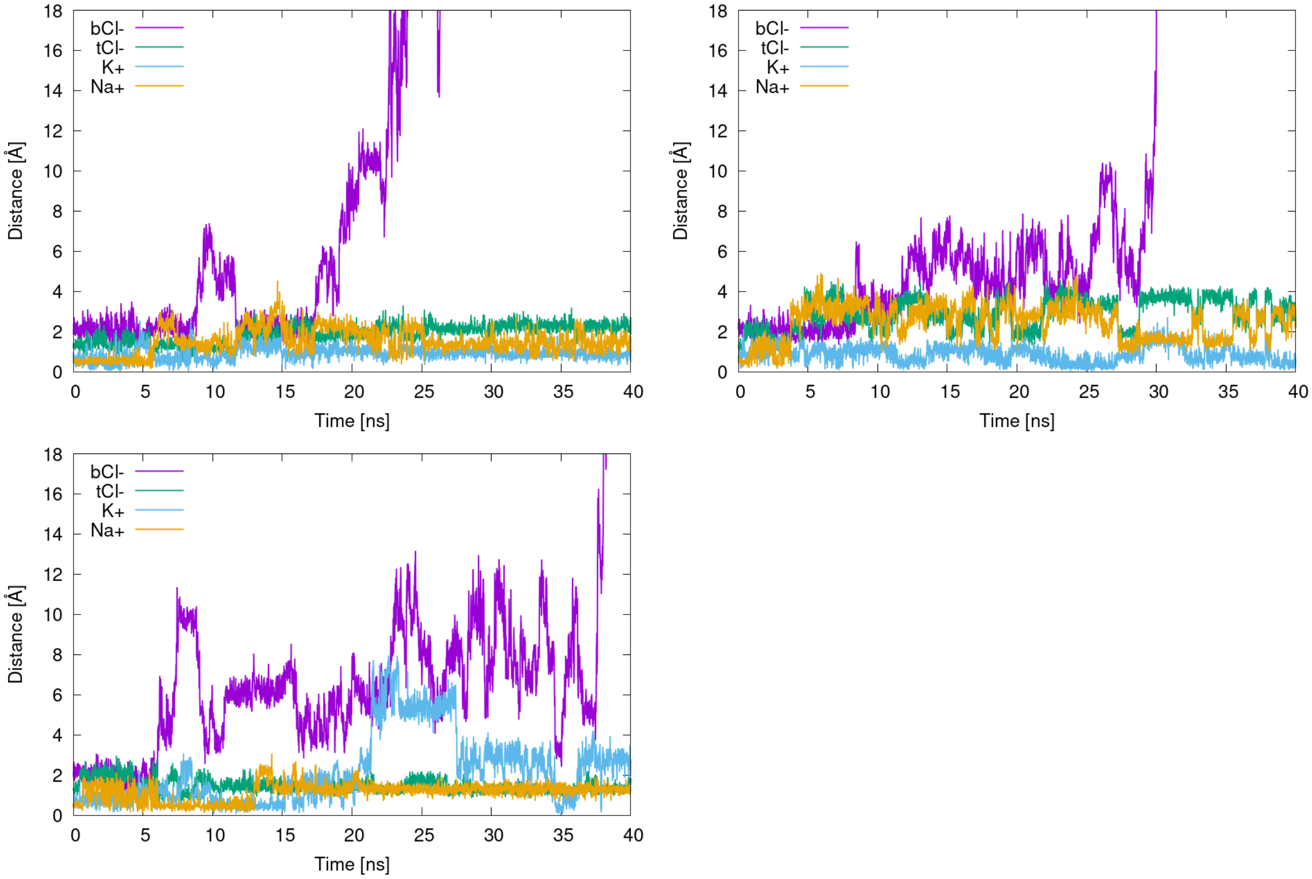

**Figure S7.** Ion dissociation events observed in preliminary metadynamics simulations of the fully ions-loaded reduced monomeric model system in which all four ions are simultaneously biased. Distance (Å) of each ion from its initial binding site vs simulation time (ns). Purple, green, cyan and orange lines refer to the bottom (b)$Cl^-$, top (t)$Cl^-$, $K^+$ and $Na^+$ ions, respectively. Three replicas of the same metadynamics simulation are shown.

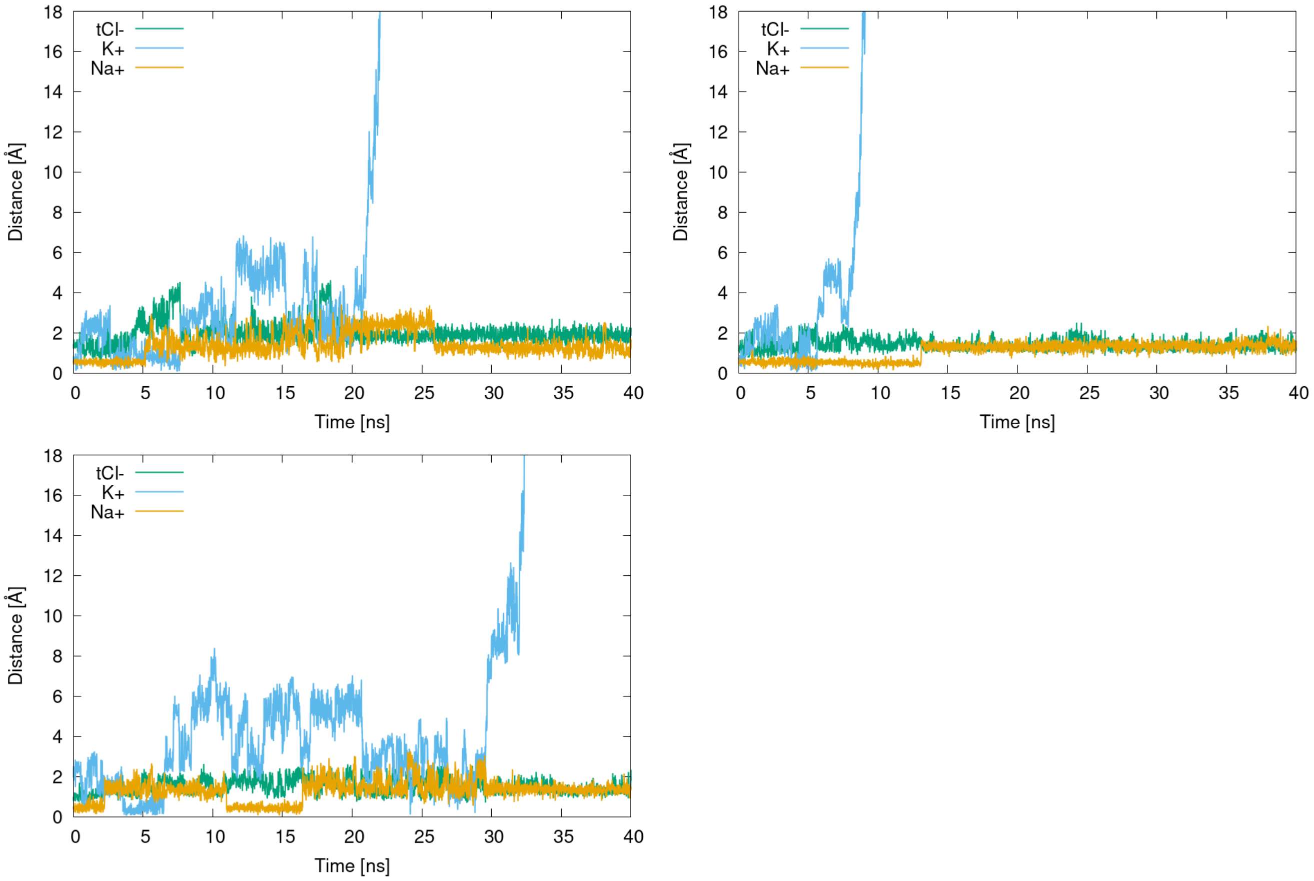

**Figure S8.** Ion dissociation events from preliminary metadynamics simulations after bCl$^-$ dissociation has occurred. Here the top (t)Cl$^-$, Na$^+$ and K$^+$ ions are simultaneously biased. Distance (Å) of the ions from their binding site vs simulation time (ns). Green, cyan and orange lines refer to tCl$^-$, K$^+$ and Na$^+$ ions, respectively. Three replicas of the same metadynamics simulation are shown.

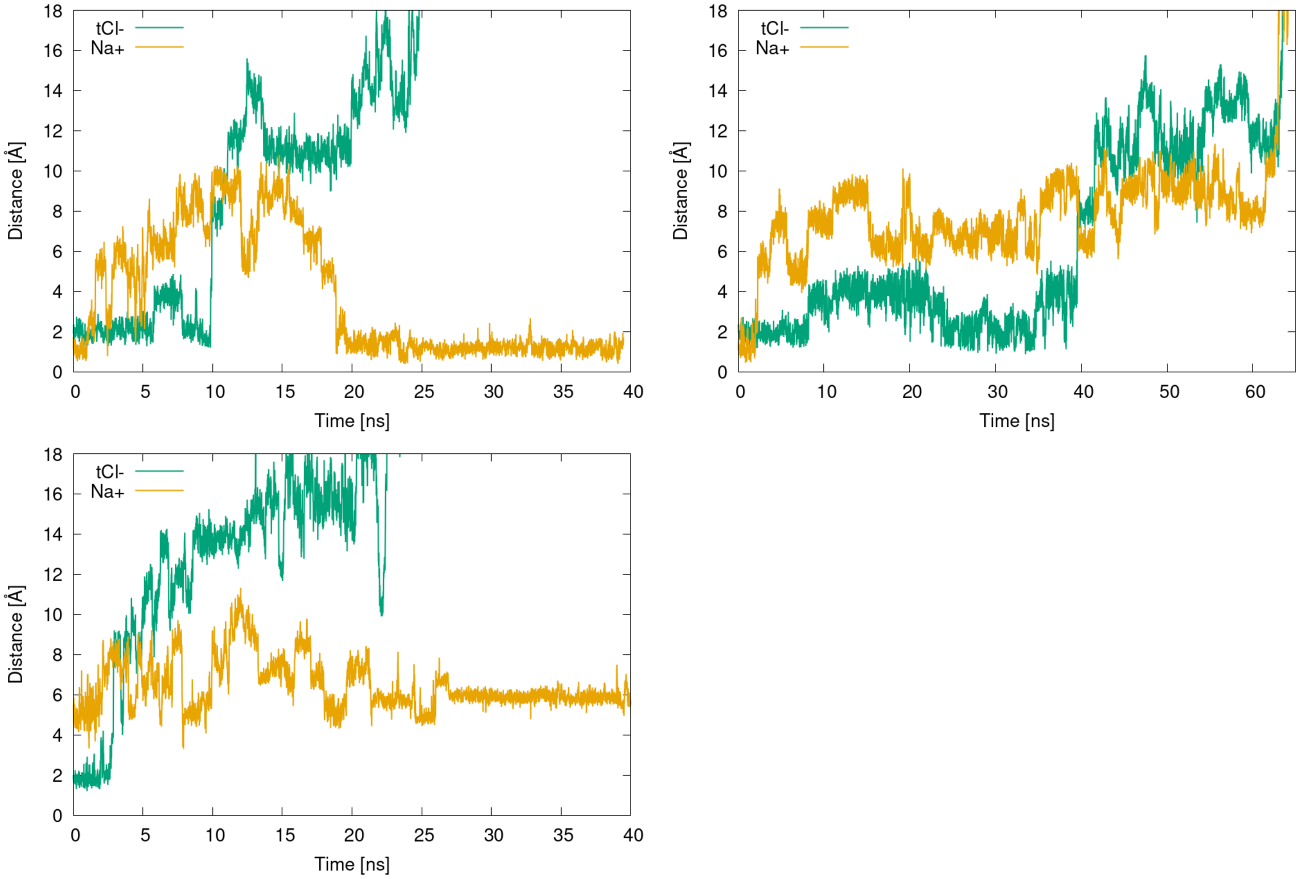

**Figure S9.** Ion dissociation events from preliminary metadynamics simulations in which the bottom $Cl^-$ and $K^+$ have already left the channel. Distance (Å) of the top (t)$Cl^-$ and $Na^+$ ions from their initial binding site vs simulation time (ns). Green and orange lines refer to the tCl- and Na+ ions, respectively.

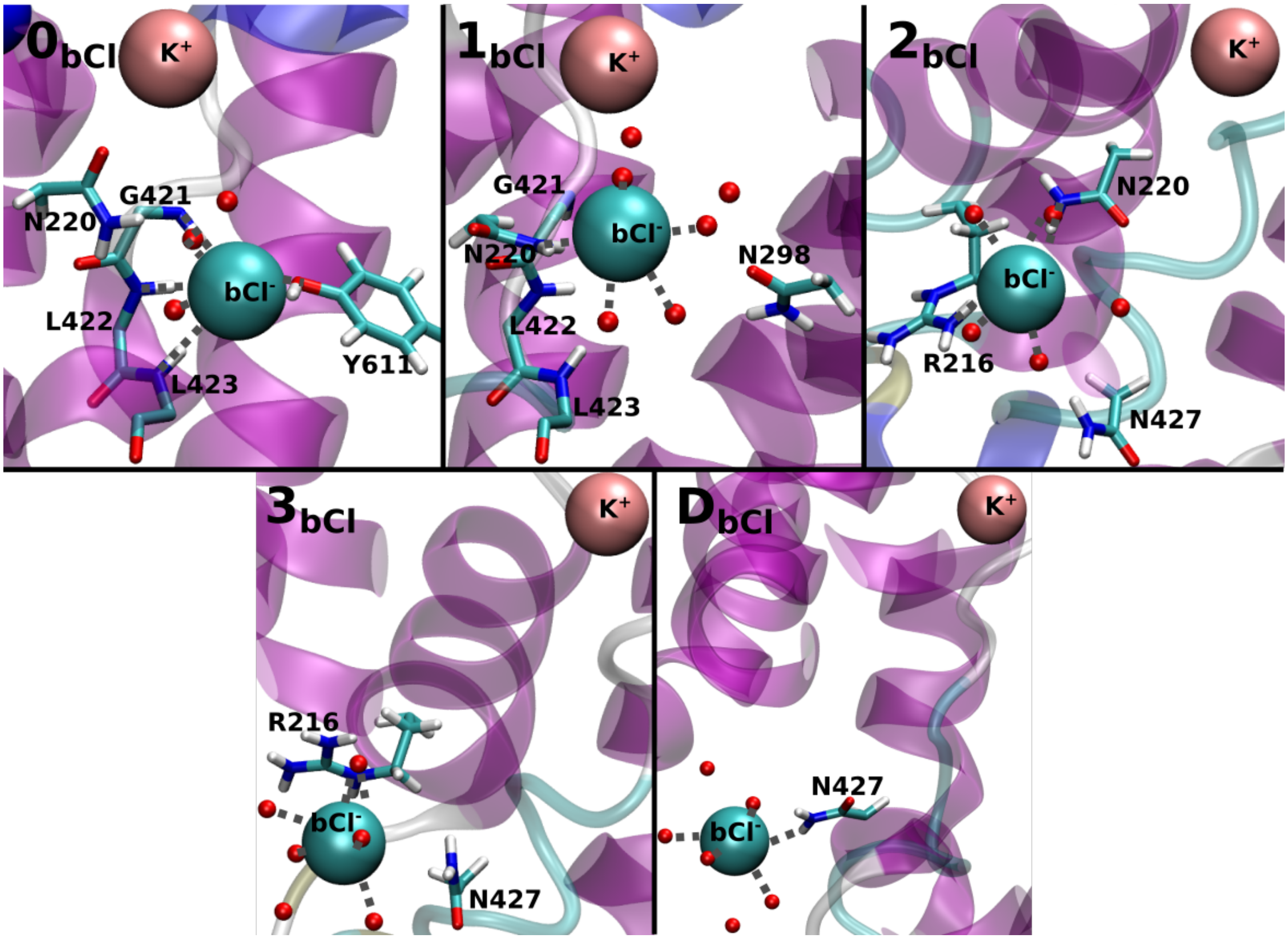

**Figure S10.** Close-up views of all minima visited by the bottom (b) $Cl^-$ ions during its translocation.

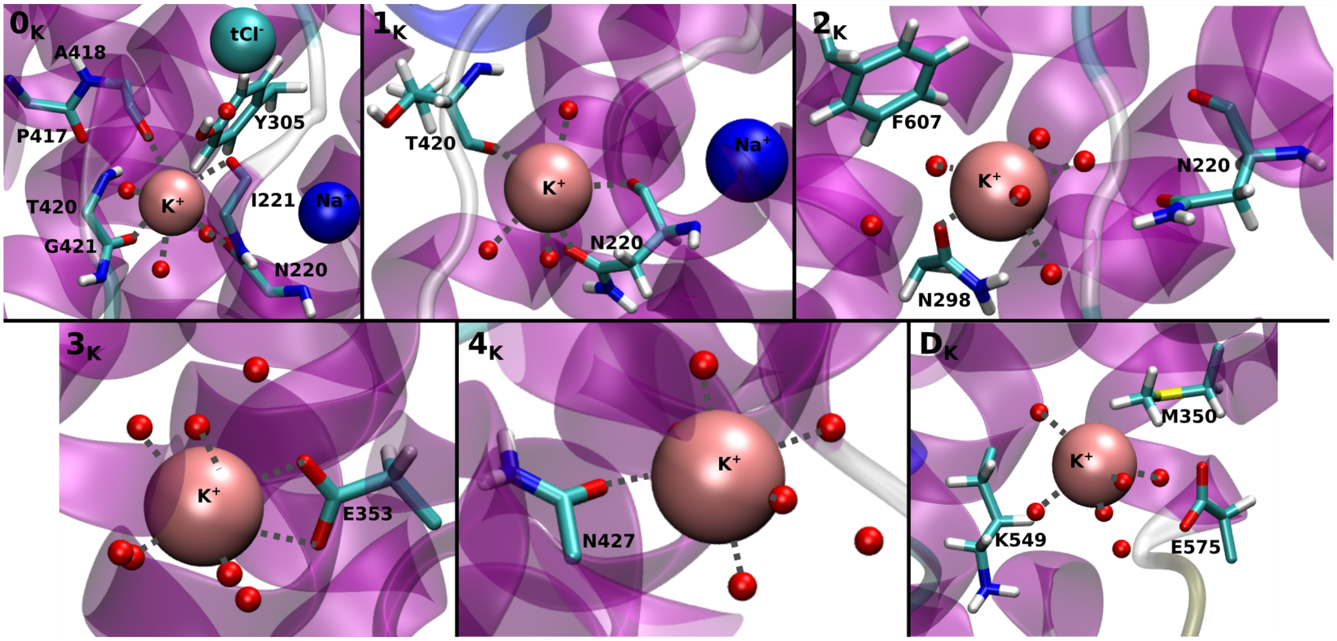

**Figure S11.** Close-up views of all the minima visited by the $K^+$ during its translocation process.

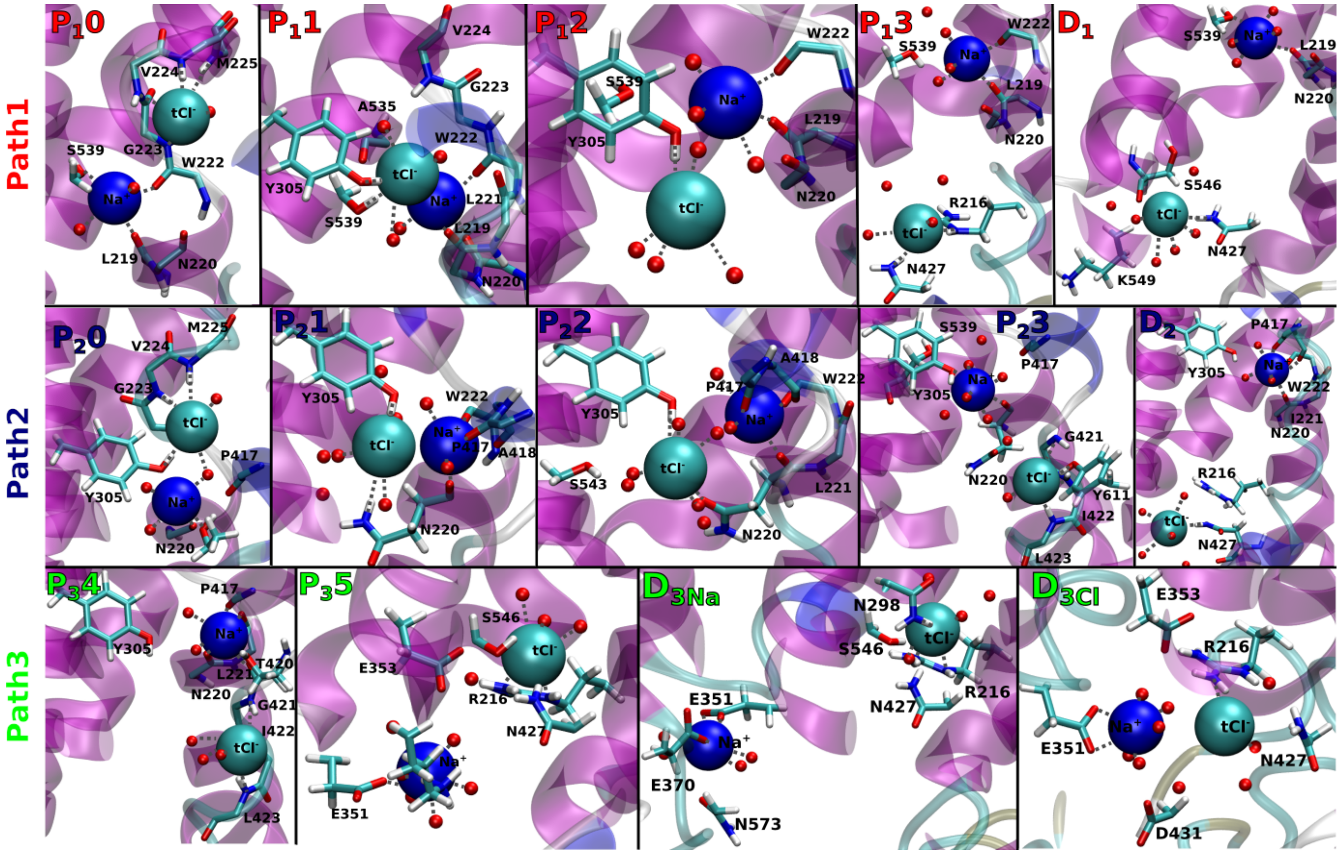

**Figure S12.** Close-up view of all the minima visited by the top(t) $Cl^-$ and $Na^+$ ions during their translocation along the different possible paths.

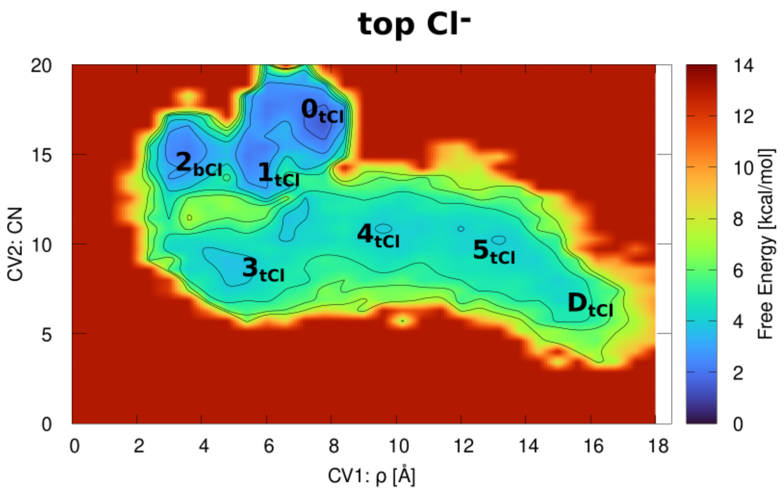

**Figure S13.** Reweighted free energy surfaces (kcal/mol) for the top (t)$Cl^-$ ion translocation reported as a function of the ion distance (Å) from protein center and of its coordination number to the protein residues in Collective Variable (CV)1 and CV2, respectively. The free energy surface (kcal/mol) isosurface is shown from blue to red color. Reweighting the $tCl^-$ dissociation onto the same set of CVs used to analyse the bottom (b)$Cl^-$ dissociation allows to better show similarities between the two $Cl^-$ translocation processes. The initial bound state minimum $\mathbf{0_{tCl}}$ and the $\mathbf{2_{bCl}}$ minimum, which corresponds to the $bCl^-$ binding site, are separated by an intermediate state ($\mathbf{1_{tCl}}$), which represents the movement of the $tCl^-$ in the vicinity of the $Na^+$ ion (as shown in the $\mathbf{P_1 1}$ and $\mathbf{P_2 1}$ states in Figure 5D of the main text). The rest of the free energy surface looks similar to the dissociation of the $bCl^-$, with intermediate state $\mathbf{3_{tCl}}$ being similar to $\mathbf{1_{bCl}}$. From this point onward the FES is rather smooth, similar to the $bCl^-$ translocation free energy surface, with two intermediate states $\mathbf{4_{tCl}}$ and $\mathbf{5_{tCl}}$ corresponding to the states $\mathbf{2_{bCl}}$ and $\mathbf{3_{bCl}}$ visited by $bCl^-$ during its dissociation.

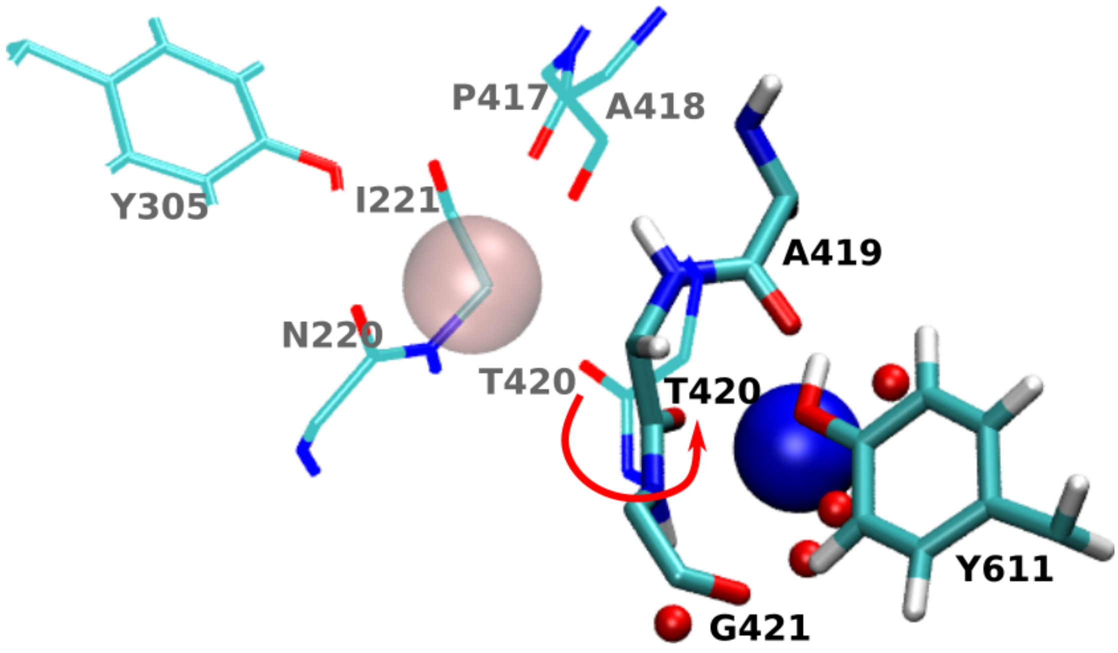

**Figure S14.** Comparison of the **-2**$_{\mathbf{Na}}$ minimum from the Na$^+$ metadynamics simulation (shown in licorice representation) and the initial K$^+$ binding site (**0**$_{\mathbf{K}}$, Figure 3 of the main text). The residues of the K$^+$ binding site are shown as lines with K$^+$ ion depicted as transparent van der Waals sphere, while the residues forming the **-2** $_{\mathbf{Na}}$ state are shown in licorice with Na$^+$ ion depicted as blue van der Waals sphere. The key difference is in the position of the Thr420 residue, which is highlighted with red arrow.

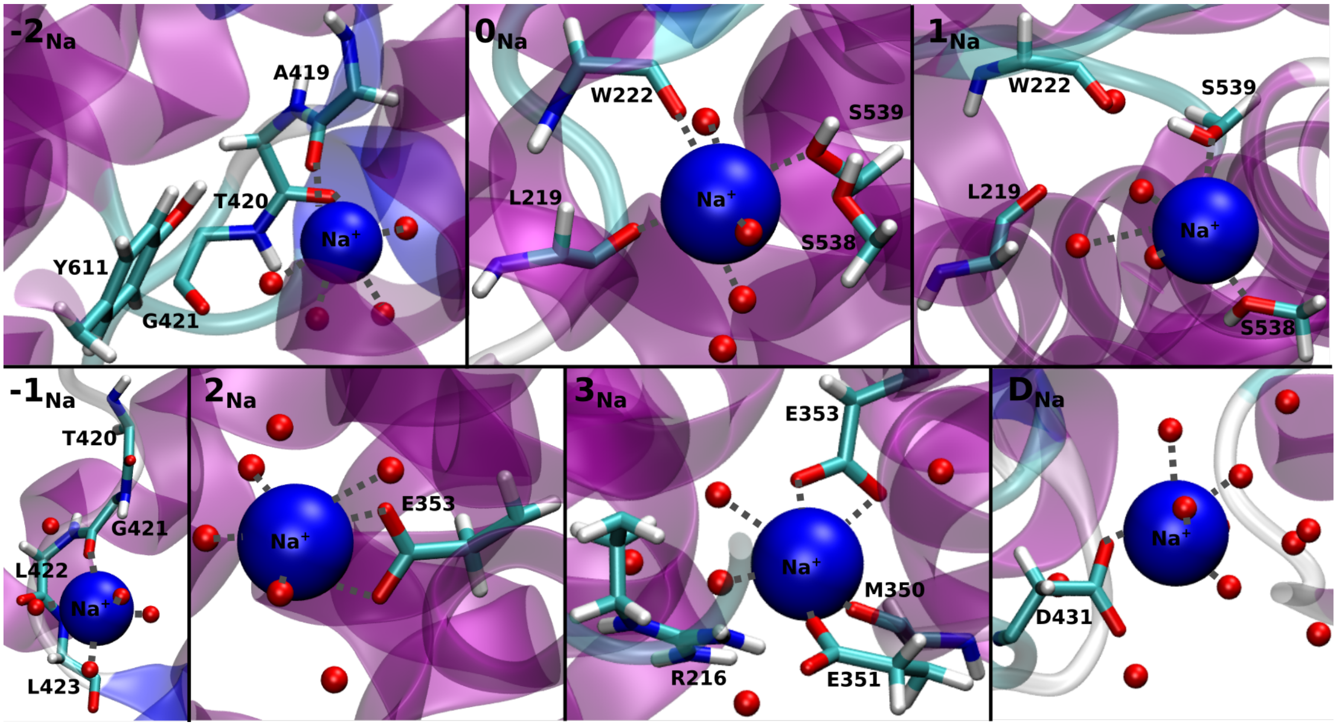

**Figure S15.** Close-ups of all the minima visited by the $Na^+$ ion during its translocation process.

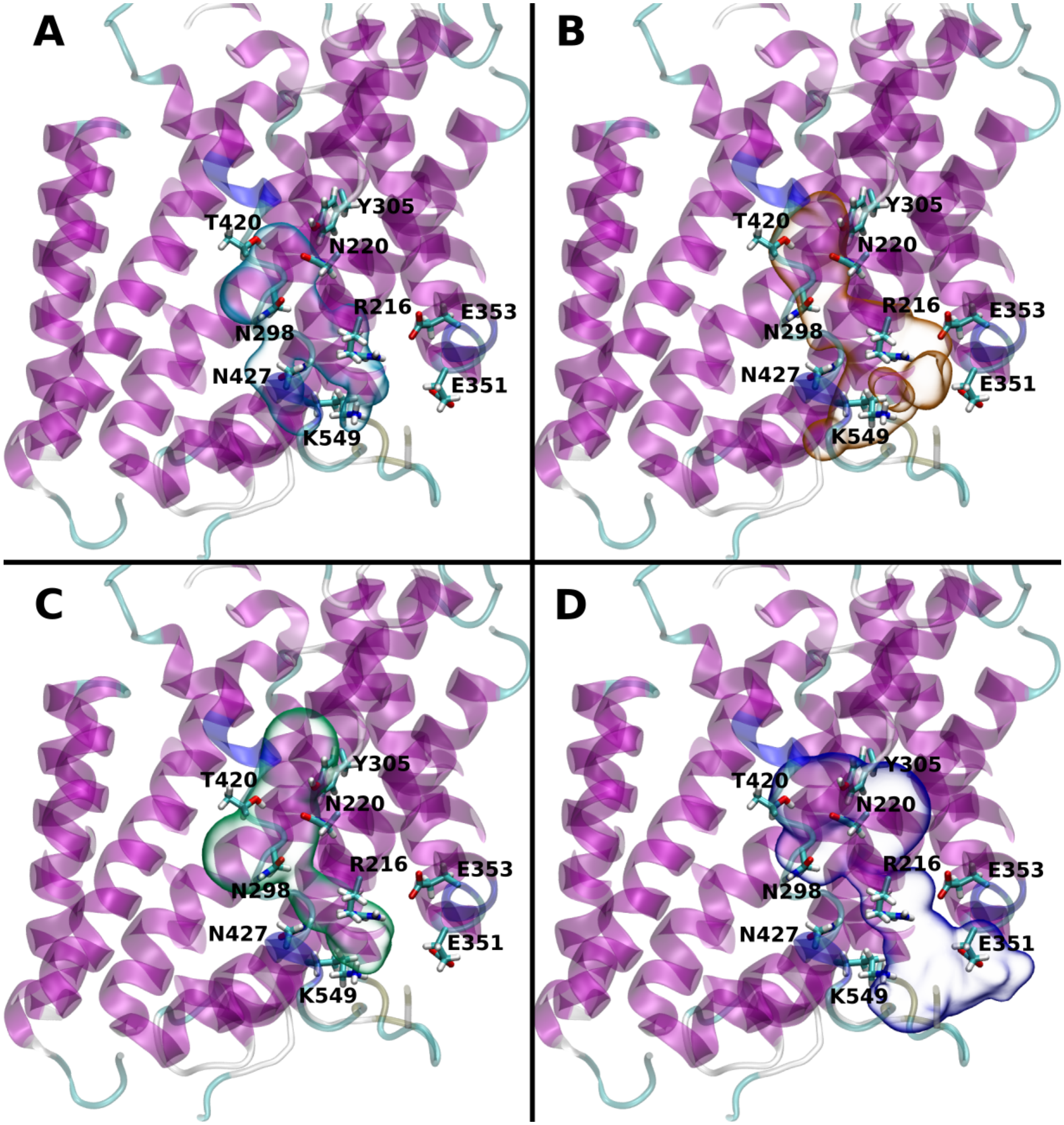

**Figure S16.** Dissociation pathways of the bCl$^-$(A), K$^+$(B), tCl$^-$(C) and Na$^+$(D) ions depicted as cyan, orange, green and blue surfaces, along the NKCC1 channel. Key residues involved in more than one ion dissociation pathway are labeled and shown as licorice. The NKCC1 protein is depicted as magenta new cartoons.

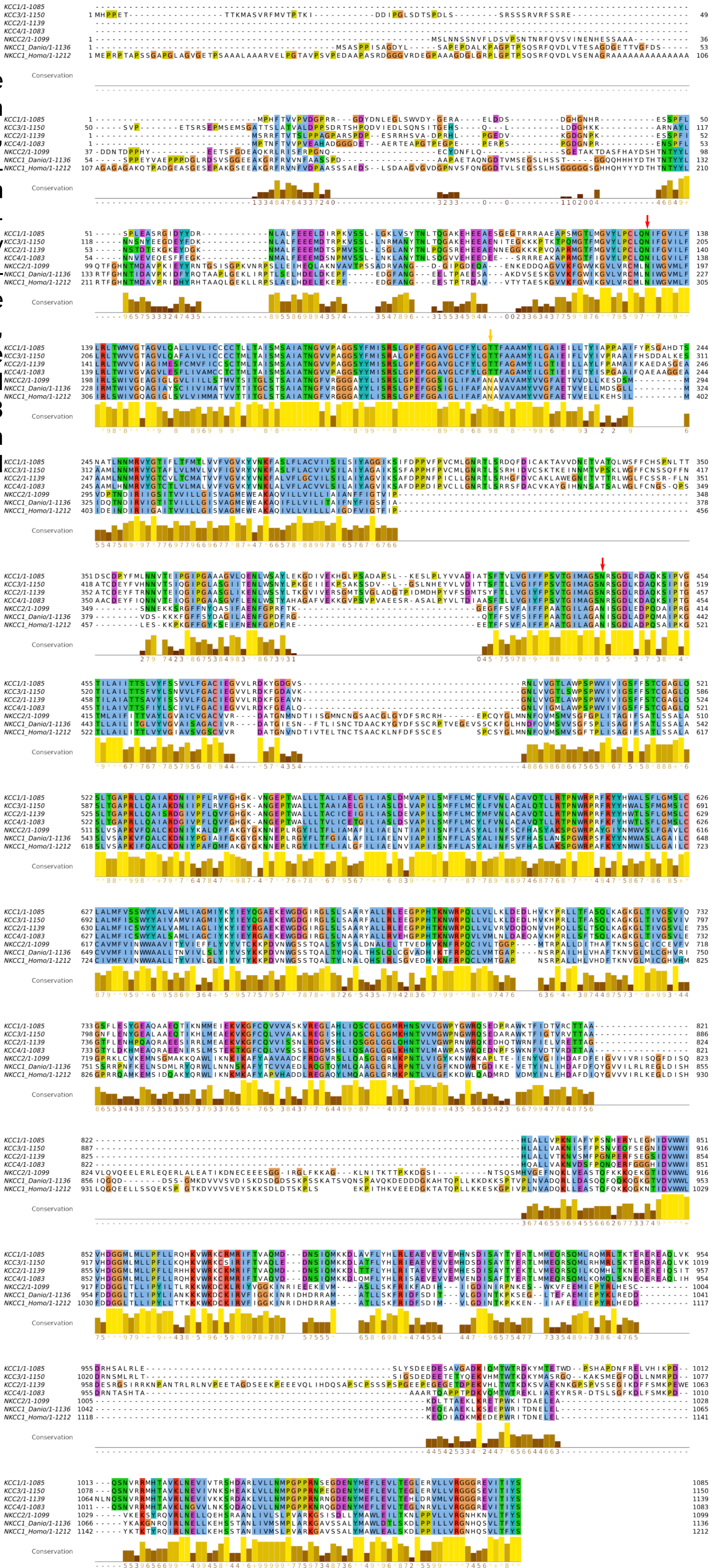

**Figure S17.** Sequence alignment of zebra fish NKCC1 (NKCC1_Danio), human NKCC1 (NKCC1_Home), human NKCC2 and human KCC1-4 ion channel generated by ClustalΩ (2, 3) and visualized by Jalview (4). The N220 and N427 residues, conserved in all of the channels, are highlighted with red arrow. The N298 residue conserved only in the NKCCs is highlighted with yellow arrow.

### Supplementary Tables

**Table S1.** List of the residues defining the initial binding sites of the four ions. Bottom (b) $Cl^-$ and top (t)$Cl^-$ refer to most cytosol-oriented and least cytosol-oriented $Cl^-$ ion, respectively. All ions exhibit a hexahedral coordination and are completed by water molecules, when six aminoacidic ligands are missing.

| Ions | Binding site |
|---|---|
| $bCl^-$ | G421@N, I422@N, L423@N, Y611@OH |
| $tCl^-$ | G223@N, V224@N, M225@N, L226@N, Y305@OH, Y454@OH |
| $K^+$ | N220@O, I221@O, Y305@OH, P417@O, T420@OG, T420@O |
| $Na^+$ | L219@O, W222@O, S538@OG, S539@OG |

**Table S2.** Overall dissociation free energy ($\Delta G^‡$) barriers (kcal/mol) for in-cell translocation of each ion. $bCl^-$ and $tCl^-$ stand for the bottom and the top $Cl^-$ ions, respectively.

| Ion | | $\Delta G^‡$ (kcal/mol) |
|---|---|---|
| $bCl^-$ | | 2.6 ± 0.7 |
| $K^+$ | | 5.8 ± 0.5 |
| $tCl^-$ | Path1 | 7.1 ± 0.8 |
| | Path2 | 6.5 ± 0.8 |
| $Na^+$ and $tCl^-$ | Path3 | 6.5 ± 0.8 |
| $Na^+$ | | 6.9 ± 0.8 |

**Table S3.** Protein residues coordinating the bottom (b)$Cl^-$ in different minima visited along its translocation pathway as observed in metadynamics simulation in which the b$Cl^-$ ion dissociation is biased. Residues in gray provide only water-mediated interactions. The ion is always hexacoordinated with water molecules completing the coordination sphere. Residues highlighted in green participate in more than one ion translocation pathway.

| $0_{bCl}$ | $1_{bCl}$ | $2_{bCl}$ | $3_{bCl}$ | $D_{bCl}$ |
|---|---|---|---|---|
| G421 | N220 | R216 | R216 | N427 |
| L422 | N298 | N220 | N427 | |
| L423 | | N427 | | |
| Y611 | | | | |

**Table S4.** Protein residues coordinating $K^+$ in different minima visited along its translocation pathway as observed in metadynamics simulation in which the $K^+$ ion dissociation is biased. Residues in gray provide only water-mediated interactions. The ion is always hexacoordinated with water molecules completing the coordination sphere. Residues highlighted in green participate in more than one ion translocation pathway.

| $0_K$ | $1_K$ | $2_K$ | $3_K$ | $4_K$ | $D_K$ |
|---|---|---|---|---|---|
| N220 | N220 | N298 | E353 | N427 | E575 |
| L221 | T420 | N220 | | | |
| P417 | P417 | | | | |
| A418 | | | | | |
| Y305 | | | | | |
| T420 | | | | | |

**Table S5.** Protein residues coordinating top (t)$Cl^-$ in different minima visited along its translocation pathway as obtained from the metadynamics simulation in which the t$Cl^-$ and $Na^+$ ions are simultaneously biased. Residues in gray provide only water-mediated interactions. The ion is always hexacoordinated with water molecules completing the coordination sphere. Residues highlighted in green participate in more than one ion translocation pathway.

| $P_1 0$ | $P_1 1$ | $P_1 2$ | $P_1 3$ | $D_1$ |
|---|---|---|---|---|
| G223 | Y305 | Y305 | R216 | N427 |

| | | | | |
|---|---|---|---|---|
| V224 | S539 | | N427 | K549 |
| M225 | | | | |
| L226 | | | | |
| **$P_20$** | **$P_21$** | **$P_22$** | **$P_23$** | **$D_2$** |
| G223 | N220 | N220 | G421 | N427 |
| V224 | Y305 | Y305 | L422 | |
| M225 | | S543 | L423 | |
| Y305 | | | Y611 | |
| **$P_34$** | **$P_35$** | **$D_{3Na}$** | **$D_{3Cl}$** | **$D_{3NaCl}$** |
| G421 | R216 | R216 | R216 | R216 |
| L422 | N427 | N427 | N427 | |
| L423 | S546 | S546 | | |
| Y611 | | N298 | | |

**Table S6.** Protein residues coordinating $Na^+$ in different minima visited along its translocation pathway as obtained from the metadynamics simulation in which the top (t)$Cl^-$ and $Na^+$ ions are biased simultaneously. Residues in gray provide only water-mediated interactions. The ions are always hexacoordinated with water molecules completing their coordination sphere. Residues highlighted in green participate in more thaen one ion translocation pathway.

| **$P_10$** | **$P_11$** | **$P_12$** | **$P_13$** | **$D_1$** |
|---|---|---|---|---|
| L219 | L219 | L219 | L219 | L219 |
| W222 | W222 | N220 | N220 | N220 |
| S539 | S539 | W222 | W222 | W222 |
| | | S539 | S539 | S539 |
| **$P_20$** | **$P_21$** | **$P_22$** | **$P_23$** | **$D_2$** |
| N220 | W222 | N220 | N220 | N220 |
| Y305 | P417 | W222 | Y305 | W222 |
| P417 | A418 | Y305 | P417 | Y305 |

| | | | | |
|---|---|---|---|---|
| T420 | | P417 | T420 | P417 |
| | | A418 | S539 | A418 |

| **$P_3$4** | **$P_3$5** | **$D_{3Na}$** | **$D_{3Cl}$** | **$D_{3NaCl}$** |
|---|---|---|---|---|
| N220 | E351 | E351 | E351 | E351 |
| L221 | E353 | E575 | D431 | |
| Y305 | | | | |
| P417 | | | | |
| T420 | | | | |

**Table S7.** Protein residues coordinating $Na^+$ in different minima visited along its translocation pathway as obtained from the metadynamics simulation in which the $Na^+$ dissociation is biased. Residues in gray provide only water-mediated interactions. Ion is always hexacoordinated with water molecules completing the coordination sphere. Residues highlighted in green participate in more than one ion translocation pathway.

| **$-2_{Na}$** | **$0_{Na}$** | **$1_{Na}$** | **$-1_{Na}$** | **$2_{Na}$** | **$3_{Na}$** | **$D_{Na}$** |
|---|---|---|---|---|---|---|
| T420 | W222 | S538 | G421 | E353 | E351 | D431 |
| A419 | L219 | S539 | L422 | | E353 | |
| | S538 | W222 | | | M350 | |
| | S539 | L219 | | | | |

### References

1. T. A. Chew, *et al.*, Structure and mechanism of the cation–chloride cotransporter NKCC1. *Nature* **572**, 488–492 (2019).
2. F. Sievers, D. G. Higgins, Clustal Omega for making accurate alignments of many protein sequences. *Protein Science* **27**, 135–145 (2018).
3. F. Madeira, *et al.*, The EMBL-EBI search and sequence analysis tools APIs in 2019. *Nucleic acids research* **47**, W636–W641 (2019).
4. A. M. Waterhouse, J. B. Procter, D. M. Martin, M. Clamp, G. J. Barton, Jalview Version 2 —a multiple sequence alignment editor and analysis workbench. *Bioinformatics* **25**, 1189–1191 (2009)., 3495–3499 (2019).
